# 5’ extended protein-coding *INO80E* transcript regulates expression of two head-to-head overlapping genes: *INO80E* and *HIRIP3*

**DOI:** 10.1101/2023.12.04.569901

**Authors:** Natalia Ryczek, Aneta Łyś, Elżbieta Wanowska, Joanna Kozłowska-Masłoń, Izabela Makałowska

## Abstract

The *INO80E* gene encodes the protein involved in the chromatin remodeling processes as a part of the multi-subunit INO80-chromatin remodeling complex. The *INO80E* gene is located on chromosome 16 and overlaps head-to-head with the *HIRIP3* gene encoding protein, which binds H2B and H3 core histones and HIRA protein and regulates the chromatin and histone metabolism. Antisense transcription of head-to-head overlapping PC genes may have several consequences and none of them have been comprehensively investigated and explained. Here, we determined that INO80E-201, which overlaps the *HIRIP3* gene transcripts, forms an R-loop at its 5’ end. We also demonstrated that overlapping transcripts of INO80E and *HIRIP3* form an RNA:RNA duplex that has the stabilizing effect on the involved mRNAs. Our results confirmed that this stabilization could be mediated by the ELAVL1 protein. We additionally introduced *de novo* methylation using the CRISPR/Cas system into the promoter sequence of *INO80E* gene. As a result of the introduced changes, reduced expression of *HIRIP3* and *INO80E* gene transcripts was observed. It was determined that methylated cytosines were located in the binding sites for four transcription factors including RARG, which was further confirmed to be important in the transcription of both studied genes. Our results strongly suggest that the formation of an RNA:RNA duplex is necessary for stable simultaneous expression of both genes. Lack of this dsRNA structure results in a loss of a wider DNA opening and in consequence transcriptional interference. We also concluded that forming R-loops probably plays only a supplementary role and is not required for proper expression of *HIRIP3* and *INO80E* gene transcripts.

## Introduction

The *INO80E* gene encodes the protein involved in the chromatin remodeling processes as a part of the multi-subunit INO80-chromatin remodeling complex (Bao and Shen 2007; Clapier and Cairns 2009; S. Zhang et al. 2016). This complex is involved in crucial cell biology processes, such as double-strand breaks repair (Poli, Gasser, and Papamichos-Chronakis 2017), cell cycle, apoptosis (Yoo et al. 2022), and chromatin accessibility during meiotic sex chromosome inactivation (Chakraborty and Magnuson 2023). It regulates the transcription of genes involved in these processes. The INO80 complex is widely described as an ATP-dependent nucleosome remodeler (Hsieh et al. 2022; Poli, Gasser, and Papamichos-Chronakis 2017), however it prefers to mobilize hexasomes over nucleosomes and thus regulate replication, transcription, and DNA repair (Hsieh et al. 2022). This complex is of great importance for the biological functions of the cell, therefore the processes which secure the expression of genes by encoding their subunits, including INO80E, are of very high interest. The *INO80E* gene is located on chromosome 16 and overlaps head-to-head with the *HIRIP3* gene encoding protein, which binds H2B and H3 core histones and HIRA protein, in order to regulate the chromatin and histone metabolism (Assrir et al. 2007; Lorain et al. 2023; Siddaway et al. 2022). Such a genomic arrangement of two genes that are crucial for proper cell functioning is quite interesting since, as many reports suggest, it may have many negative effects.

Overlaps between protein-coding (PC) genes, i.e. partial or complete sharing of genomic location, were identified in prokaryotes as well as in eukaryotes, including humans (Bøvre and Szybalski 1969; Henikoff et al. 1986; Makałowska, Lin, and Hernandez 2007; X.-J. Wang, Gaasterland, and Chua 2005). There are several types of possible genes overlaps: the 5’-ends (head-to-head) overlap, the nested genes, the 3’-end (tail-to-tail) overlap, and genes overlapping with their promoter regions, of which the implications of gene overlap at the 5’ ends are some of the least studied (Rosikiewicz, Suzuki, and Makałowska 2018; Shearwin, Callen, and Egan 2005; Wight and Werner 2013). Antisense transcription of head-to-head overlapping PC genes may have several consequences and none of them have been comprehensively investigated and explained. One of the most reported is the potential occurrence of transcriptional interference, which can cause aberrations in the expression of overlapping genes (Rosikiewicz and Makałowska 2016). However, works published on this subject are in a big part theoretical and do not clarify this phenomenon (Callen, Shearwin, and Egan 2004; Chia et al. 2017; Osato et al. 2007; Shearwin, Callen, and Egan 2005). Nevertheless, it is quite plausible that two head-to-head overlapping genes are expressed from bidirectional promoters. Competition for this promoter may result in weaker transcription (Trinklein et al. 2004). Similar effect may have other possible types of transcriptional interference, such as polymerase collision, the so called “sitting duck” interference or occlusion (Shearwin, Callen, and Egan 2005). The study performed by Bendtsen et al. shows that RNA polymerase (RNAP), by binding to one promoter, can repress the activity of the overlapping promoter located on the opposite DNA strand (Bendtsen et al. 2011).

Transcripts of overlapping genes may regulate transcription at the RNA–DNA and RNA-RNA interaction levels. Examples of the first one could be DNA methylation and demethylation or downregulation of the expression of the sense gene by antisense RNA (Cebrat et al. 2008). Chromatin modification and silencing of the sense promoter have also been demonstrated (Yu et al. 2008). Another way of expression regulation at the RNA-DNA interaction level is the formation of R-loops. The term R-loop refers to the DNA-RNA hybrid structure and the displaced single-stranded DNA (ssDNA) (Aguilera and García-Muse 2012). Regardless of being for a long time described as a rare byproduct of transcription, there exist examples demonstrating that R-loops naturally formed during transcription can play a beneficial role in transcription regulation at certain loci (Richard and Manley 2017). Advances in technology and an increasing number of studies have shown that R-loops facilitate physiological processes, and in certain contexts, they can also cause genome instability (Crossley, Bocek, and Cimprich 2019; García-Muse and Aguilera 2019; Richard and Manley 2017). Recent studies have shown that head-to-head transcription promotes the formation of an R-loop structure, which has been associated with positive expression regulation when two genes are in close proximity on opposite DNA strands (Boque-Sastre et al. 2015; Tan-Wong, Dhir, and Proudfoot 2019). Nevertheless, regulatory effects of R-loops formation were confirmed only for overlapping pairs of protein-coding genes and long non-coding RNAs (lncRNAs), for example, the *RASSF1* gene and its antisense lncRNA *ANRASSF1* (*RASSF1* antisense RNA 1) or antisense lncRNA *VIM-AS1* (VIM antisense RNA 1), which overlaps the vimentin (*VIM*) gene at the 5’ end (Beckedorff et al. 2013). So far, published results demonstrate that R-loop formation could be associated with a positive regulation in the expression of two transcripts overlapping in a head-to-head manner.

The complementarity of two antisense transcripts may lead to RNA–RNA interactions by the formation of RNA duplexes (Su et al. 2012). Well known are duplexes between protein-coding mRNAs and lncRNAs (Nojima and Proudfoot 2022; Statello et al. 2020) and there are many reports explaining their regulatory function (Herman, Tsitsipatis, and Gorospe 2022; Nojima and Proudfoot 2022; Statello et al. 2020). As shown by Hastings et al., such a duplex may physically hide access to splicing sites, which may result in the formation of alternative splice forms (Hastings et al. 1997). RNA:RNA duplexes may also have an influence on transcript transport (Khochbin and Lawrence 1989), contribute to endo-siRNA (Werner et al. 2014), or have a stabilizing effect on protein-coding sense transcripts by blocking the RNA destabilizing motif (Uchida et al. 2004) or by masking microRNA target sites (Faghihi et al. 2010). Also, the formation of duplexes may be a mechanism enabling the unwinding of the DNA strands, increasing the access of RNA Pol II to promoters, and, consequently, increasing transcription. The phenomenon of RNA duplexes formation from two complementary fragments of protein-coding transcripts has not yet been studied, although, according to Sharma et al., the presence of the mRNA interactions at the 5’ ends is highly abundant (Sharma et al. 2016). The occurrence of the RNA duplex formed by two mRNAs and its functions have been so far described only for the head-to-head overlapping genes *TP53* and *WRAP53* (Mahmoudi et al. 2009).

Overlap at the 5’ ends between two PC genes may be either fixed or intermittently present. This mostly depends on the usage of alternative transcription start sites (TSS). Some gene pairs may have TSSs that will always cause overlap with the gene on the opposite strand of the DNA (C. H. Chen, Pan, and Lin 2019; Rosikiewicz, Suzuki, and Makałowska 2018). There are also genes that use multiple TSSs for transcription - proximal and more distal. When they use the distal TSS/TSSs, there is an overlap with the gene on the other strand of the DNA, however, when they use only proximal TSS/TSSs, the overlap does not occur (Rosikiewicz et al. 2021). This dynamically changes under the influence of various intra- and extra-cellular conditions (Makhnovskii et al. 2022; Rosikiewicz et al. 2021; Tatip et al. 2020). It has been described that changes in the TSSs used by certain genes have a translation regulatory function, e.g., in cancers, such as squamous cell carcinomas (Weber et al. 2022) and during cerebellar development (P. Zhang et al. 2017). More precisely, Weber and colleagues described an increased overall translation efficiency of gene isoforms with extended 5’UTRs, as well as the enriched presence of ribosomal proteins and splicing factors on these isoforms (Weber et al. 2022). TSSs switching has also been observed and described for protein-coding genes and antisense lncRNAs, for example in breast cancer (Maruyama et al. 2012) and neural differentiation (Ahmad et al. 2017).

Despite many studies, the exact function of overlapping transcripts and their role in expression regulation are unknown. Most research results and hypotheses suggest that head-to-head PC gene overlap phenomenon has a downregulating effect on involved genes’ expression. Chen et al. for example, in support of antisense regulation by forming dsRNA, showed that human sense–antisense transcripts tend to be co-expressed and/or inversely expressed more frequently than expected by chance (J. Chen et al. 2005). Henz et al. also observed negatively correlated expression of overlapping transcripts (Henz et al. 2007). Our previous studies, however, did not demonstrate any negative effects due to gene overlap. In contrast, genes utilizing overlapping TSSs had, on average, higher expression levels than the same genes utilizing only non-overlapping TSSs (Rosikiewicz et al. 2021). Therefore, the head-to-head overlap of *INO80E* and *HIRIP3* genes raises important questions on how their expression is regulated and what is the role of overlapping transcripts. This is particularly important because both genes are crucial for the proper functioning of cells. Thus, in our research the primary focus lied upon elucidation of the mechanism regulating the expression of the *INO80E* and *HIRIP3* genes, while simultaneously providing empirical evidence that interactions between the 5’ ends of two PC transcripts is essential for proper transcription.

## Results

### *INO80E* and *HIRIP3* genomic organization and expression

According to the Ensembl (Martin et al. 2023) and NCBI (Sayers et al. 2022) databases, the *INO80E* gene has fourteen transcriptional forms, one of which is elongated at the 5’ end (INO80E-201). The *HIRIP3* gene has seven transcriptional forms, from which four (HIRIP3-204, HIRIP3-202, HIRIP3-206, and HIRIP3-207) overlap with the INO80E-201 transcript and only two, HIRIP3-201 and HIRIP3-204, are annotated as protein-coding (Figure 1A). We confirmed in HEK293T cells the presence of all *HIRIP3* transcripts and five *INO80E* transcripts, which were selected for further studies and presented in Figure 1A, including INO80E-201 (Figure 1B).

**Figure 1.**
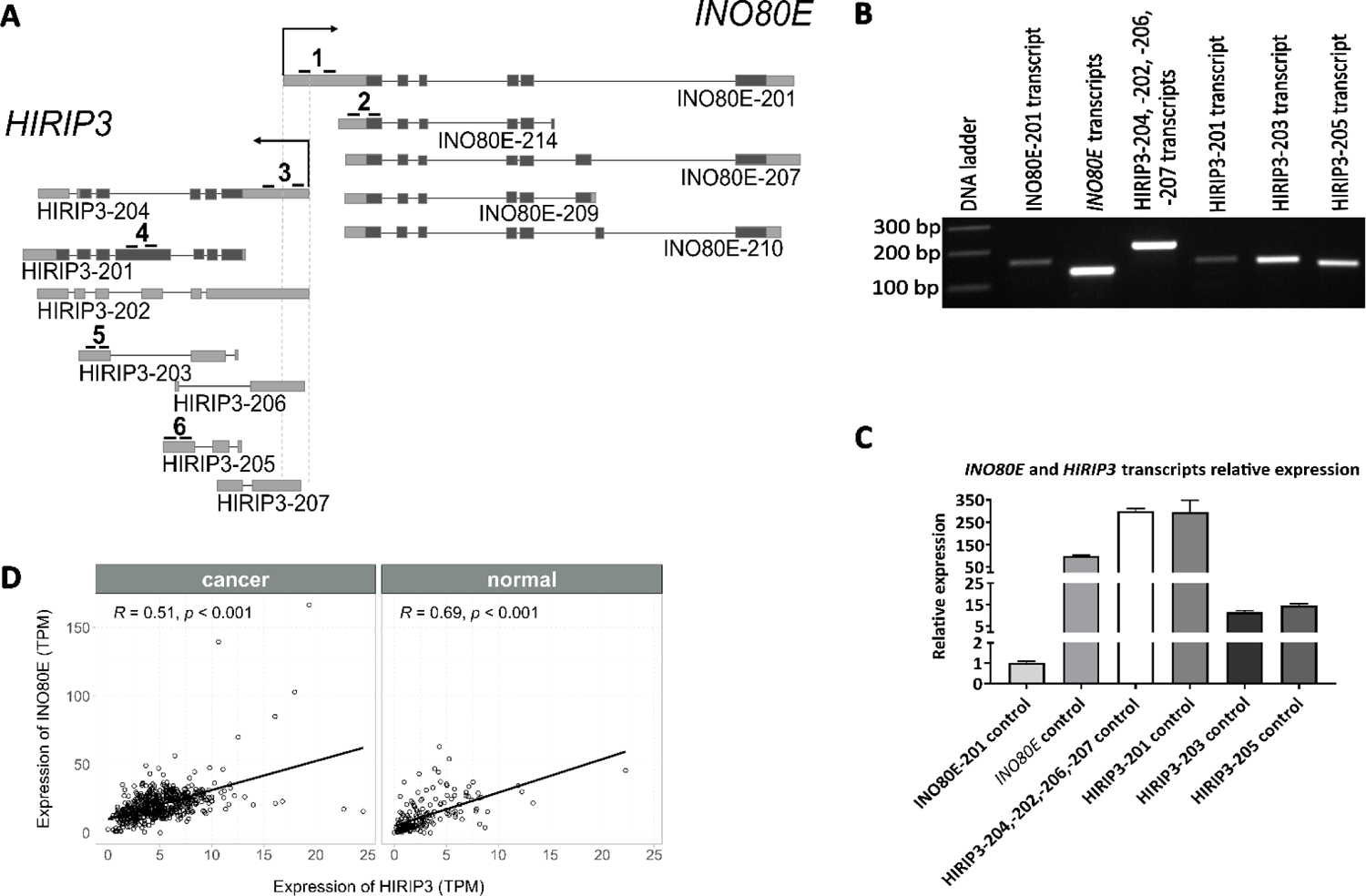
**A.** Transcripts of the *HIRIP3* and selected transcripts of the *INO80E* gene pair expressed in HEK293T cells. The HIRIP3-204, HIRIP3-202, HIRIP3-206, HIRIP3-207, and INO80E-201 transcripts overlap at the 5’ ends. Light grey represents non-coding exon sequences, while dark grey –coding exon sequences. Numbers represent primers used for amplification of: 1 – INO80E-201 transcript; 2 – *INO80E* transcripts; 3 – HIRIP3-204, HIRIP3-202, HIRIP3-206, and HIRIP3-207 transcripts; 4 – HIRIP3-201 transcript; 5 – HIRIP3-203 transcript; 6 – HIRIP3-205 transcript. **B.** *HIRIP3* and *INO80E* transcripts expression in HEK293T cells. **C.** Relative expression of *HIRIP3* and *INO80E* transcripts in HEK293T cells. **D.** Correlation between *INO80E* gene and *HIRIP3* gene expression (TPM) in cancer and non-cancer cells.

Following the confirmation of transcripts expression, we estimated and compared their abundance. Overall, in HEK293T cells the *HIRIP3* gene is expressed at a higher level than the *INO80E* gene. In the case of *INO80E* the expression was estimated separately for the transcript extended at the 5’ end and for remaining isoforms. Due to a bigger variation in transcripts structures and transcription start sites, *HIRIP3* transcript expression was examined in four groups. Transcripts overlapping with the INO80E-201 transcript (HIRIP3-204, HIRIP3-202, HIRIP3-206, and HIRIP3-207) were analyzed together. The remaining three transcripts were examined separately (Figure 1C). Next, we determined that the quantitative ratio of INO80E-201 transcript to other transcripts of *INO80E* gene is 1:766. This very low level of overlapping INO80E-201 may suggest a regulatory role rather than protein coding. The quantitative ratio of *HIRIP3* transcript expression was also calculated. The expression of overlapping transcripts was also lower in comparison with other HIRIP3 transcripts, but the difference was not as large as in the case of *INO80E*.

It has been suggested that, due to possible transcriptional interference, overlapping genes have a negative expression correlation (J. Chen et al. 2005; Henz et al. 2007). Utilizing the results of our previous studies of 784 RNA-seq samples encompassing diverse tissues and cell lines (Szcześniak et al. 2020), we performed an analysis of the expression correlation of these genes in non-cancerous and cancerous tissues/cell lines samples. In both cases, a strong and statistically significant positive correlation was observed (Figure 1D). Similarly, a positive expression correlation was found when data was divided according to the tumor type and the organ system (Supplementary materials, Figure S1A and S1B). These results confirm our previous report, demonstrating a positive correlation of expression of 5’ overlapping genes (Rosikiewicz et al. 2021).

The positive correlation of *INO80E* and *HIRIP3* gene expression allowed us to hypothesize the existence of some co-regulatory mechanisms. To investigate the mutual influence of *INO80E* and *HIRIP3* on their expression pattern, we performed several knock-down experiments with the Antisense LNA GapmeRs (Qiagen) followed by qPCR. First, we silenced all transcripts of the *HIRIP3* gene. The results showed us that after *HIRIP3* gene silencing, the relative expression of the INO80E-201 transcript decreased to minimal levels, as the rest of the *INO80E* gene transcript expression increased (Figure 2A). These results show that indeed there is a link between the expression of these two genes. Therefore, we checked how a knock-down of the *INO80E* gene transcripts will affect expression of *HIRIP3* (Figure 2B). Under such conditions, the expression of overlapping *HIRIP3* transcripts (HIRIP3-204, HIRIP3-202, HIRIP3-206, and HIRIP3-207) decreased by almost 80%. However, there was a 10-fold increase in the expression of the HIRIP3-201 transcript. For the remaining two transcripts of the *HIRIP3* gene (HIRIP3-203 and HIRIP3-205), no significant changes were observed (Figure 2B).

**Figure 2.**
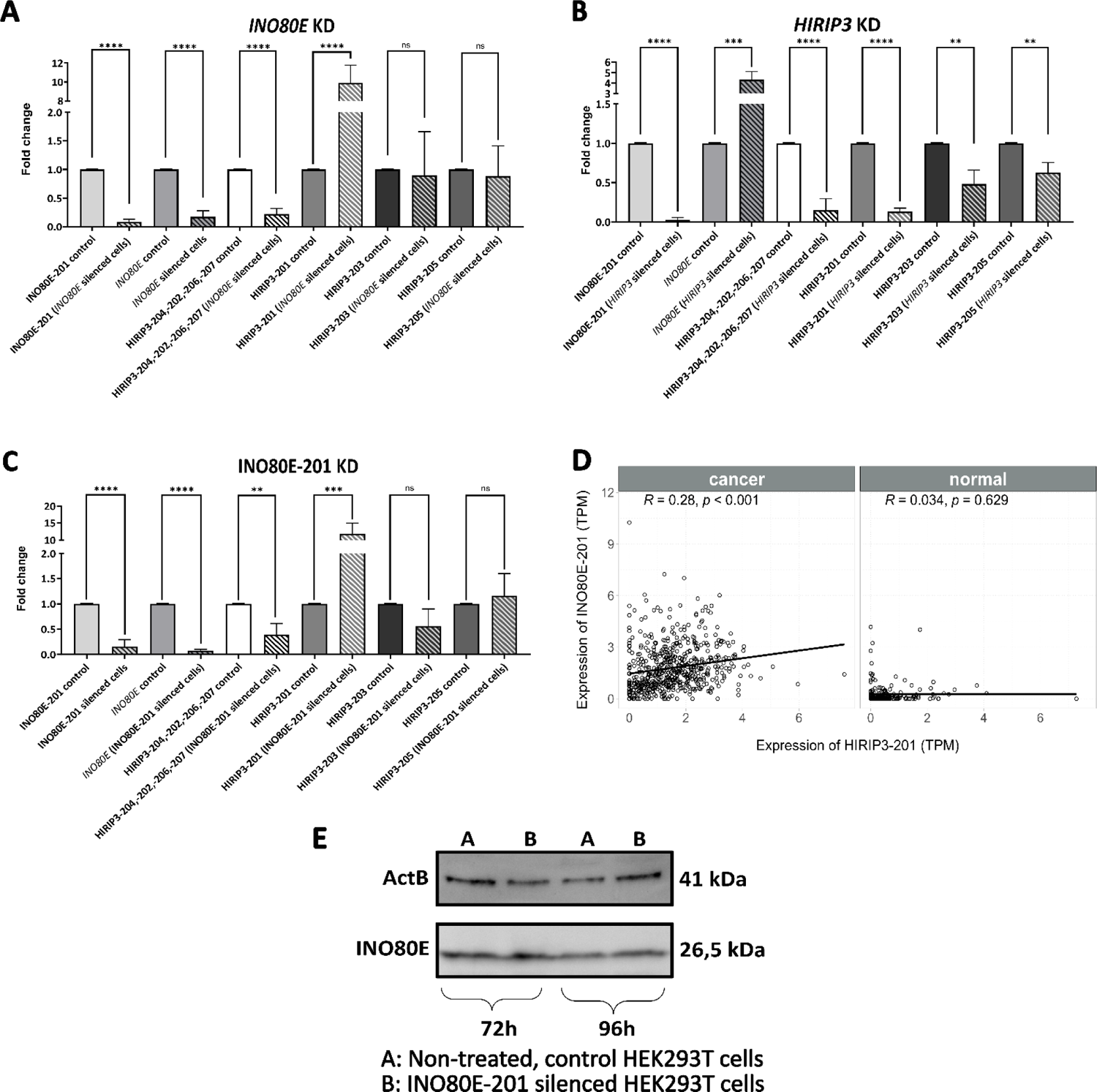
Fold change of *HIRIP3* and *INO80E* transcripts expression after silencing HIRIP3 (A), *INO80E* (B), and INO80E-201 (**C**) (ns – not significant; ** - p<0.01; *** - p<0.001; **** - p<0.0001). Barred columns represent knocked-down cells of the corresponding transcripts. **D.** Correlation between INO80E-201 and HIRIP3-201 transcripts expression (TPM) in cancer and non-cancer tissues/cells. **E.** INO80E protein level 72 h and 96 h after silencing of the INO80E-201 transcript in HEK293T cells. A: Non-treated, control HEK293T cells B: INO80E-201 silenced HEK293T cells. ActB - beta Actin Polyclonal Antibody; INO80E - INO80E Polyclonal Antibody.

To determine if the obtained results are related to the silencing of the 5’-extended transcript of the INO80E-201 gene, an additional knock-down experiment was performed (Figure 2C). After INO80E-201 was knocked-down, the expression of all *INO80E* gene transcripts decreased almost 90%. This shows that the presence of the INO80E-201 transcript is required for a higher expression of other transcripts of this gene. The silencing of INO80E-201 had also an effect on the expression of *HIRIP3* gene transcripts. An almost 70% decrease in expression of the 5’-extended *HIRIP3* gene transcripts was noticed and the expression of the HIRIP3-201 transcript increased thirteen times. No changes in HIRIP3-203 and HIRIP3-205 transcripts expression were noted (Figure 2C). All knock-down results, with tested control samples, including the *MALAT1* silencing as a positive control, Negative A GapmeR (Qiagen) treatment as negative control, and Lipofectamine 3000 treatment as a transfection control, are shown in the Supplementary Materials (Supplementary materials, Figure S2).

The silencing experiments had a significant effect on transcripts of both overlapping genes and indicate a regulatory role of INO80E-201. The quantitative ratio of INO80E-201 transcript to all transcripts of the *INO80E* gene also suggests such a function. Considering positive expression correlation of analyzed genes, as shown in Figure 1D, and an increased expression of HIRIP-201 after silencing of *INO80E,* we checked the expression correlation of these two transcripts utilizing the same RNA-seq data from normal and cancerous tissues. It turned out that the expressions of these transcripts are not correlated with each other (Figure 2D) in normal cells, however, there is a low but significant positive correlation in cancer tissues. Lack of correlation in normal cells may seem to be contradictory to our result, showing that silencing of INO80E-201 caused an increase of HIRIP3-201. However, this may be a result of a lack of such a relation in the opposite way, i.e., changes in HIRIP3-201 have no influence on INO80E-201.

Next, we examined whether a silencing of the INO80E-201 transcript, and in result a decrease of the expression level of the *INO80E* gene as a whole, influences the INO80E protein level in HEK293T cells. Western blot analysis after 72h and 96h post-transfection showed that there were no significant differences in the amount of the INO80E protein (26.5 kDa – isoform 1, UniProt: Q8NBZ0-1; 15.2 kDa – isoform 2, UniProt: Q8NBZ0-2) in control HEK293T cells versus cells with a silenced INO80E-201 (Figure 2E).

### Subcellular localization

Obtained results indicated that the main function of overlapping transcripts may lie in expression regulation. If so, they should be localized predominantly in the nucleus. To confirm this, their localization was checked by a subcellular fractionation followed by qPCR. It has been established that INO80E-201 is located mainly in the nucleus and associated with chromatin (Figure 3A), which confirms the presumption about its regulatory function. *HIRIP3* transcripts that are involved in the formation of an overlap are similarly abundant in the cytoplasm and the nucleus. HIRIP3-203 and HIRIP3-205 are located mainly in the cell nucleus. The role of these two transcripts is unknown but they were not affected by the *INO80E* gene or INO80E-201 transcript silencing. All other transcripts of the *INO80E* gene and the remaining *HIRIP3* gene transcript, HIRIP3-201, are found in the greatest amount in the cytoplasm (Figure 3A). This information is consistent with data available in the NCBI and Ensembl databases, where these transcripts are labeled as encoding proteins.

**Figure 3.**
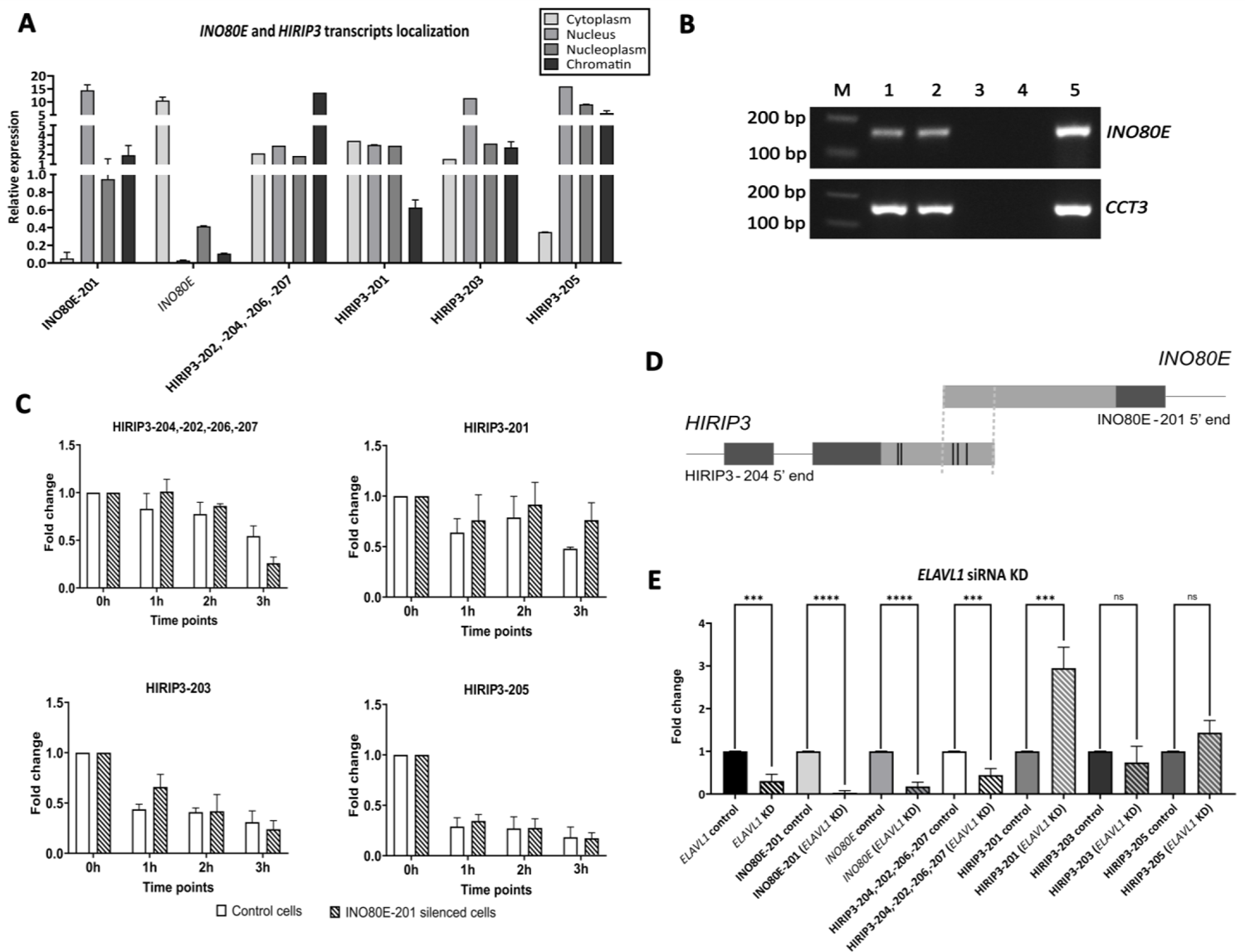
**A.** Location of *INO80E* and *HIRIP3* transcripts in control/standard conditions. **B.** The results of the RAP followed by PCR detection of captured transcripts. The 1,5% agarose gel electrophoresis results. PCR detection of *INO80E* gene was performed to confirm duplexes between *INO80E*/*HIRIP3* after RAP and CCT3 to confirm positive control duplexes between *CCT3*/*OIP5-AS1*. M - Thermo Scientific™ GeneRuler 100 bp Plus DNA Ladder (100-3000 bp), 1, 2 – PCR products on cDNA from HEK293T treated with AMT crosslinker, two biological repetitions; 3, 4 – PCR products on cDNA from HEK293T without crosslinker, two biological repetitions; 5 – control cDNA from total HEK293T RNA; 6 – control of the purity of the PCR reaction. **C.** Degradation of *HIRIP3* gene transcripts in HEK293T cells in control/standard conditions and after silencing the INO80E-201 transcript. Barred columns represent INO80E-201 knock-down cells (ns – not significant; *-p<0.05; ***-p<0.001; **** - p<0.0001). **D.** ELAVL1 protein binding motifs predicted in 5’UTR sequence of the *HIRIP3* gene (marked in black vertical lines). **E.** *ELAVL1* gene knock-down effect on the *INO80E* and *HIRIP3* genes (ns – not significant; *-p<0.05; **-p<0.01; **** - p<0.0001). Barred columns represent *ELAVL1* knock-down cells.

### RNA:RNA duplexes and transcripts stability

Focusing on the regulatory role of overlapping transcripts associated with chromatin, we checked whether the complementary 5’UTR fragments of involved transcripts interact with each other and form RNA:RNA duplexes. The possibility of forming RNA:RNA duplexes by transcripts of protein-coding genes has already been confirmed, but has been mostly based on bioinformatics analyses of high-throughput sequencing data (Lu, Gong, and Zhang 2018; Sharma et al. 2016; M. Zhang et al. 2021). The only verified example comes from studies of overlapping *WRAP53* and *TP53* genes (Mahmoudi et al. 2009). We checked whether *INO80E* and *HIRIP3* are involved in the formation of the RNA:RNA duplex through the RNA Antisense Purification assay, followed by PCR detection (Engreitz, Lander, and Guttman 2015). The 4’-aminomethyl trioxsalen (AMT) fixed RNA duplexes were captured using a biotinylated oligonucleotide probe complementary to the transcripts of the *HIRIP3* gene (Supplementary Materials, Table S1). During PCR detection, transcripts of *INO80E* were identified. This indicates the occurrence of RNA duplexes between *INO80E* and *HIRIP3* genes (Figure 3B).

The formation of RNA:RNA duplexes between transcripts of the *INO80E* and *HIRIP3* genes may have several consequences, such as: inhibition of splicing or mRNA transport, recruitment of various proteins, and induction of mRNA instability (Kumar and Carmichael 1998). The complementary joining of transcripts may also have a stabilizing function and could delay mRNA degradation. Therefore, we tested whether the formation of the *INO80E*:*HIRIP3* RNA duplex contributes to an increased stability of the *HIRIP3* gene transcripts. To check this, two types of cells were prepared: control/untreated cells and cells after 72h transfection with the Antisense LNA GapmeRs (Qiagen) aimed at the INO80E-201 transcript. After treatment with actinomycin D (ActD), the expression of *HIRIP3* gene transcripts was checked at three time points (1h, 2h, 3h) and compared to cells not treated with ActD. This experiment showed that upon silencing of INO80E-201, the overlapping *HIRIP3* gene transcripts, HIRIP3-204, HIRIP3-202, HIRIP3-206, and HIRIP3-207, degrade faster than under controlled conditions without INO80E-201 silencing. The effect was seen three hours after the addition of ActD (Figure 3C). This indicates the stabilizing function of the RNA duplex in the cell nucleus. We also observed slower degradation of the HIRIP3-201 transcript upon silencing of INO80E-201, which confirms other results showing an increased expression of this transcript under these conditions. For the other tested *HIRIP3* gene transcripts, no changes were observed (Figure 3C).

In addition to the mRNA stabilizing function, RNA:RNA duplexes can provide a scaffold for the proteins necessary for the correct expression and processing of overlapping genes in the cell nucleus. Therefore, utilizing RAP-MS analysis, we checked which proteins bind to the transcripts involved in pairing. The HEK293T cells were UV-crosslinked and subjected to an RNA Antisense Purification procedure using biotinylated probes (Table S1). Captured proteins were then used for MS analysis. This experiment revealed two main groups of proteins: related to R-loops and those involved in splicing and mRNA stability (Table 1).

**Table 1.** Proteins associated with *INO80E*:*HIRIP3* RNA duplex. * - R-loop related proteins; # - proteins related to R-loop and involved in splicing and mRNA stability; without mark - proteins involved in splicing and mRNA stability.

| Protein symbol | Protein full name | Score |
| --- | --- | --- |
| TADA2B | Transcriptional adapter 2-beta | 1479 |
| CNOT9 | CCR4-NOT transcription complex subunit 9 | 516 |
| NPM1 * | Nucleophosmin | 409 |
| HNRNPC | Heterogeneous nuclear ribonucleoproteins C1/C2 | 407 |
| HNRNPA1 | Heterogeneous nuclear ribonucleoprotein A1 | 267 |
| LSM4 | LSM4 homolog, U6 small nuclear RNA and mRNA degradation associated | 251 |
| HNRNPH1 | Heterogeneous nuclear ribonucleoprotein H1 | 214 |
| HNRNPA2B1 | Heterogeneous nuclear ribonucleoproteins A2/B1 | 189 |
| HNRNPK | Heterogeneous nuclear ribonucleoprotein K | 98 |
| SSBP1 * | Single stranded DNA binding protein | 96 |
| HNRNPH3 | Heterogeneous nuclear ribonucleoprotein H3 | 93 |
| PARP1 * | Poly [ADP ribose] polymerase 1 | 88 |
| HNRNPU | Heterogeneous nuclear ribonucleoprotein U | 79 |
| KHSRP # | KH type splicing regulatory protein | 72 |
| HNRNPF | Heterogeneous nuclear ribonucleoprotein F | 71 |
| ELAVL1 | ELAV like RNA binding protein 1 | 66 |
| SON | SON DNA and RNA binding protein | 57 |
| HNRNPM # | Heterogeneous nuclear ribonucleoprotein M | 56 |
| TARDBP # | TAR DNA binding protein 43 | 44 |
| DDX39A # | ATP dependent RNA helicase DDX39A (DEXD box helicase 39A) | 43 |
| SNRPF | Small nuclear ribonucleoprotein F | 42 |
| HNRNPA3 | Heterogeneous nuclear ribonucleoprotein A3 | 42 |
| NONO # | Non POU domain containing octamer binding protein | 41 |
| SNRPD2 | Small nuclear ribonucleoprotein Sm D2 | 41 |
| PCBP2 | Poly(rC)-binding protein 2 | 37 |

The INO80E-201 transcript, forming an RNA:RNA duplex with the overlapping *HIRIP3* gene transcripts, may act by blocking the formation of the secondary mRNA structure at the 5’ ends, thus revealing the binding sites of proteins with a stabilizing function. Such a protein, ELAV Like RNA Binding Protein 1 - ELAVL1, has been identified in RAP-MS as binding to captured transcripts. ELAVL1 selectively binds to AU-rich regions that signal degradation of mRNA (Bakheet et al. 2018; Ma, Chung, and Furneaux 1997). Using the Scan for Motifs bioinformatics tool (Biswas and Brown 2014) and RNA-Binding Protein DataBase (RBPDB) (Cook et al. 2011), we confirmed the presence of ELAVL1 protein binding motifs in the 5’UTR sequence of the *HIRIP3* gene. The sequences containing this motif are involved in the formation of the RNA:RNA duplex but also are located right next to it (Figure 3D). To check whether this protein affects the stability of *HIRIP3* and *INO80E* transcripts, siRNA silencing of the *ELAVL1* gene was performed. We found that when the *ELAVL1* gene is silenced, expression of all *INO80E* gene transcripts decreases. The expression of *HIRIP3* transcripts that are overlapping with the *INO80E* gene also decreases. The expression of the HIRIP3-201 transcript significantly increased, which is in line with silencing experiment which also demonstrated an increased expression of HIRIP3-201 when INO80E-201 was decreased (Figure 3E, Supplementary materials - Figure S3). Altogether, this experiment shows that RNA:RNA duplexes between *INO80E* and *HIRIP3* are stabilized by the ELAVL1 protein.

### R-loop formation

The presence of proteins related to the regulation of R-loops in RAP-MS gave rise to the hypothesis that R-loops may take part in the regulation of the expression of the investigated genes. There are many reports which show both the negative and positive effects of the formation of R-loops on gene expression. Examples of negative effects are RNA polymerase II pausing and genome instability due to inducement of DNA double-strand breaks (Castillo-Guzman and Chédin 2021; Gan et al. 2011; Hegazy, Fernando, and Tran 2020; Kaneko et al. 2007; Stirling et al. 2012; Wimberly et al. 2013). R-loops may positively affect expression level through epigenetic alterations, such as histone acetylation (P. B. Chen et al. 2015; Ginno et al. 2013; Hegazy, Fernando, and Tran 2020; Sanz et al. 2016). The possibility of R-loops formation within the studied pair of genes, *INO80E* and *HIRIP3*, was first checked in the R-loopBase database (Lin et al. 2022). Analyzes showed the presence of an R-loop in the 5’UTR region of the INO80E-201 transcript. The in vivo presence of these structures was then confirmed by a native bisulfite sequencing method (Figure 4A, Supplementary materials - Figure S4) (Boque-Sastre, Soler, and Guil 2017).

**Figure 4.**
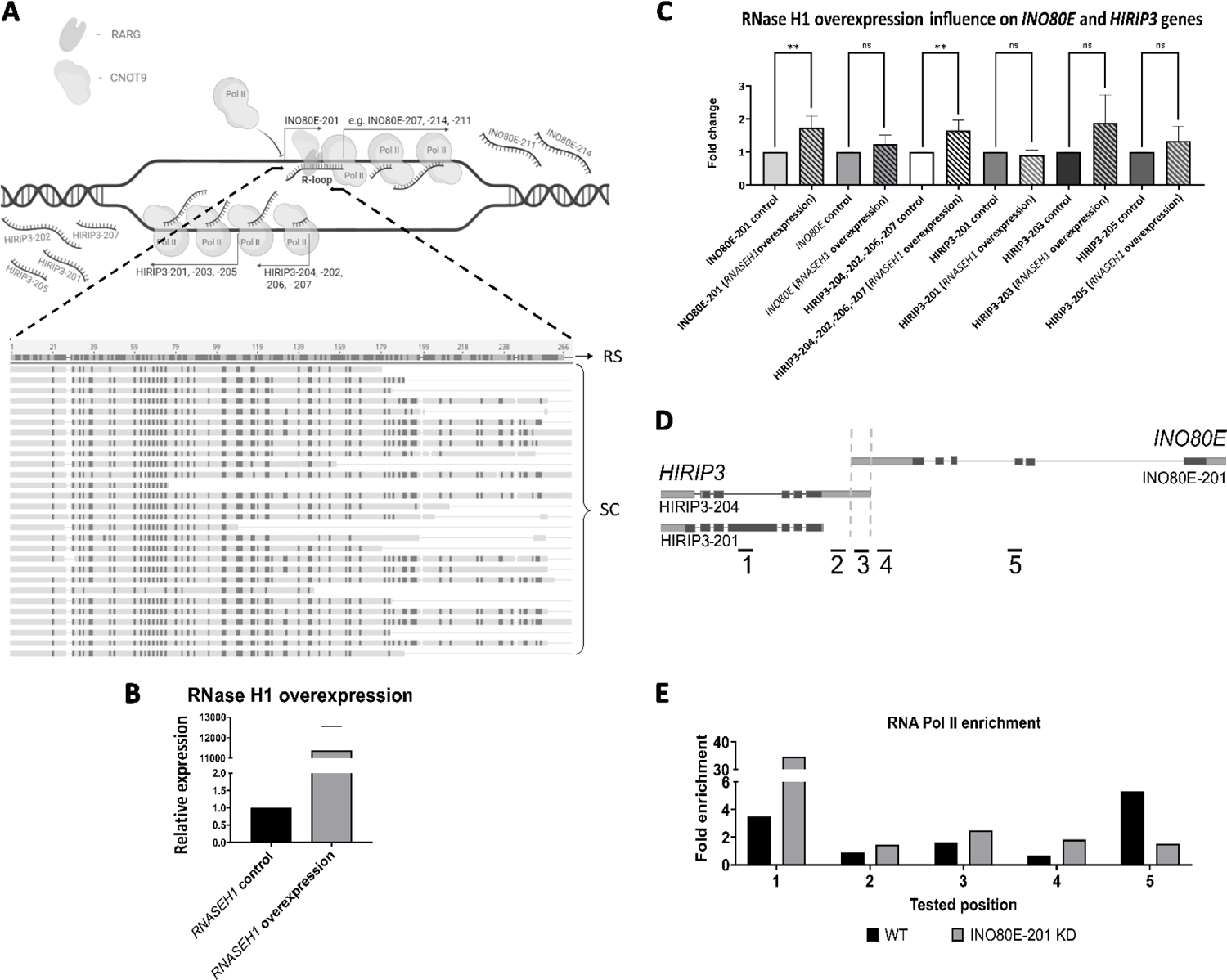
**A.** R-loop formation within the *INO80E* gene transcript INO80E-201 and results from native bisulfite sequencing R-loop method. The sequenced clones (SC) are compared to the INO80E-201 encoding sequence (RS – reference sequence). **B.** RNase H1 overexpression confirmation in HEK239 cells after FACS. **C.** RNase H1 overexpression influence on *INO80E* and *HIRIP3* genes expression. Barred columns represent knock-down cells of the corresponding transcripts. **D.** Location of the RNA Polymerase II binding detection primers for chromatin immunoprecipitation with an RNA Polymerase II antibody. **E.** Chromatin immunoprecipitation with an RNA Polymerase II antibody (ab10338, Abcam). The numbers indicate the location of the RNA Polymerase II binding detection primers.

The presence of an R-loop within the 5’UTR of the overlapping transcript may cause RNA polymerase II pausing and block transcription of INO80E-201. It may also promote transcription of remaining INO80E isoforms and *HIRIP3* without any danger of transcriptional interference. To test this hypothesis, we performed an experiment with human RNase H1 overexpression. RNase H1 unravels existing R-loops and therefore its overexpression causes a significant R-loop decrease. If the R-loop formed at the INO80E-201 5’ end plays an important regulatory role, we should observe changes in the expression of *HIRIP3* and *INO80E* transcripts. Firstly, the HEK293T cells were transfected with a pEGFP-RNASEH1 plasmid. The positive effect of the performed transfection was checked using a microscope with fluorescence detection (Supplementary materials - Figure S7). Next, cells were subjected to fluorescence-activated cell sorting (FACS). GFP-expressing cells were collected and the effectiveness of the RNase H1 overexpression was checked with qPCR. The level of RNase H1 increased over 11 500 times compared to control/untreated cells (Figure 4B). Overexpression of RNase H1 resulted in a significantly increased expression of the INO80E-201 transcript involved in the R-loop formation and overlapping *HIRIP3* transcripts. For the non-overlapping *INO80E* gene transcripts, HIRIP3-203, HIRIP3-205, and HIRIP3-201, no significant changes in the expression were noted (Figure 4C). This suggests that R-loop formation blocks the transcription of INO80E-201, which in turn leads to a faster degradation of overlapping HIRIP3 transcripts. This, however, has no effect on not overlapping transcripts of both genes.

An observed increase in the expression of the INO80E-201 transcript can be expected since R-loops block polymerase binding to the promoter. A forming R-loop may also prevent the formation of RNA:RNA duplexes and in consequence faster degradation of overlapping HIRIP3 transcripts. This would explain the increased expression of these transcripts when RNase H1 was overexpressed. However, the lack of an effect on not overlapping transcripts of *HIRIP3* was a little bit surprising, as many reports indicated a positive effect of R-loops on genes located on opposite DNA strand (Boque-Sastre et al. 2015; Rondón and Aguilera 2019).

### Transcriptional interference

The results obtained so far indicate that for transcription regulation of INO80E and HIRIP3, forming RNA:RNA duplexes is more important than R-loop formation. We hypothesize that these duplexes help to keep the chromatin more open in order to eliminate or at least decrease the possibility of transcriptional interference. This could explain a lower expression level of both genes after INO80E-201 silencing. This hypothesis was tested by a polymerase activity study utilizing a ChIP experiment with an RNA Polymerase II antibody. ChIP was performed on wild type cells and cells 72h after INO80E-201 silencing (Supplementary materials - Figure S5). The results of the experiment revealed a fold enrichment of RNA Polymerase II in the promoter region of the *HIRIP3* and *INO80E* genes after INO80E-201 silencing.

In the body of *INO80E,* significant diminishment of RNA Polymerase II was observed (Figure 4D-E). In the region of the HIRIP3-201 transcript however, an enrichment of Polymerase II RNA was detected, which supports previous results which demonstrated an increased expression of HIRIP3-201 upon INO80E-201 silencing. These results may indicate that a lack of the RNA:RNA duplex due to INO80E-201 silencing results in polymerase collision in the promoter region and as a result a weaker transcription of non-overlapping *INO80E* transcripts and overlapping *HIRIP3* transcripts. In addition, overlapping *HIRIP3* transcripts degrade faster due to a lack of stabilizing duplexes. This in turn makes upstream located TSS of HIRIP3-201 more accessible for polymerase and results in an increased expression.

### *INO80E* promoter *de novo* methylation

In the final step of the study of regulatory dependence of *INO80E* and *HIRIP3,* we altered both gene promoter regions. *De novo* cytosine methylations were introduced into the sequence between none overlapping *INO80E* TSSs and *HIRIP3* with dCas9-DNMT3A. Five tested cell lines showed changes in the expression of *INO80E* and *HIRIP3*, however, the amount of *de novo* introduced cytosine methylations was different in each line (Supplementary materials - Figure S6). For further analysis we selected cell lines No. 6 and 9, where the biggest effects of methylation were observed (Figure 5A). Significant expression decrease was noticed in all transcripts except HIRIP3-205, where no changes were noted, and HIRIP-203, who’s expression slightly decreased in cell line 6. The observed decrease in the expression of HIRIP3-201 could indicate some common regulatory elements at the DNA level.

**Figure 5.**
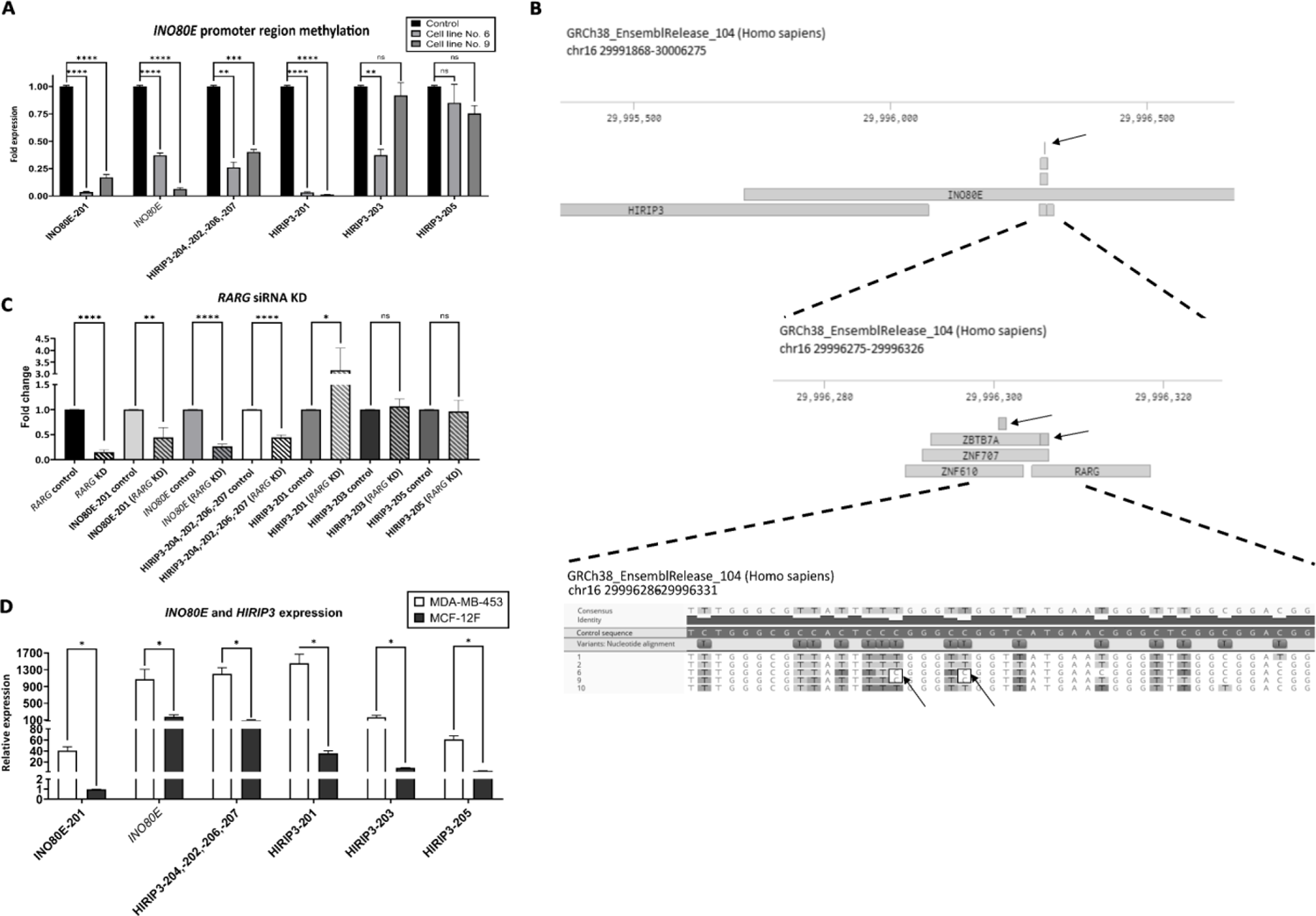
**A.** The fold expression of *INO80E* and *HIRIP3* transcripts after promoter methylation in two selected HEK239 cell lines (ns – not significant; **-p<0.01; **** - p<0.0001). **B.** Genomic localization of site-specific methylations identified in cell lines 6 and 9. 1, 2, 6, 9, 10 – numbers of tested cell lines, RARG, ZNF610, ZNF707 and ZBTB7A – target sites for these transcription factors. **C**. RARG gene knock-down effect on the INO80E and *HIRIP3* genes (ns – not significant; *-p<0.05; **-p<0.01; *** - p<0.001). Barred columns represent knocked-down cells of the corresponding transcripts. **D**. *INO80E* and *HIRIP3* gene expression in the breast cancer cell line MDA-MB-453 and the breast non-cancerous cell line MCF-12F (ns – not significant; *-p<0.05).

The localization of individual methylated cytosines was closely examined. Two modification sites specific only to cell lines 6 and 9 were identified (Figure 5B). Within their location, there are binding sites for four transcription factors: retinoic acid receptor gamma (RARG), zinc finger protein 610 (ZNF610), zinc finger protein 707 (ZNF707), and zinc finger and BTB domain containing 7A (ZBTB7A) (Figure 15). We selected RARG for further research, as the RAP-MS experiment identified the CNOT9 protein that interacts with the RARG protein.

To further investigate the involvement of the RARG protein in the expression regulation of the *INO80E* and *HIRIP3* gene pair, the *RARG* gene was knocked down using siRNA. The strongest RARG silencing effect was observed for transcripts: INO80E-201, HIRIP3-202, HIRIP3-204, HIRIP3-206, HIRIP3-207, and total expression of *INO80E*. The expression of these transcripts significantly decreased (Figure 5C, Supplementary materials - Figure S8). An increased expression was again observed in the case of HIRIP3-201. The obtained results show that the RARG protein is involved in the regulation of *INO80E* gene transcription. This experiment also confirms our previous results, which demonstrated that the expression of the *HIRIP3* and *INO80E* genes depends on the presence of the INO80E-201 transcript.

### Elevated expression in cancer

Both analyzed genes were previously indicated as associated with cancer (Prendergast et al. 2020; Yoo et al. 2022; S. Zhang et al. 2016). The INO80E protein, as part of the INO80 chromatin remodeling complex, takes part in oncogenic transcription (S. Zhang et al. 2016) and the unraveling of R-loops, enabling intensive DNA replication and, consequently, increasing the proliferation and viability of cancer cells (Prendergast et al. 2020; Yoo et al. 2022). The HIRIP3 protein interacts with the HIRA histone chaperone, the complex associated with the promotion of breast cancer (Assrir et al. 2007; Lorain et al. 2023). Additionally, our analysis of RNA-seq data revealed that the expression of INO80E-201 and HIRIP3-201 transcripts is positively correlated in cancer, while no correlation is observed in normal cells. Therefore, we checked whether and how their expression is altered in tumor cells, e.g. breast cancer, compared to unaffected breast cells. The results showed us that there is a significantly higher expression of all inspected transcripts in MDA-MB-453 breast cancer cells compared to the breast control cell line MCF-12F (Figure 5D).

### Regulatory mechanism

Results of the performed experiments demonstrate that the overlapping of the *HIRIP3* transcript of *INO80E* (transcript 201) plays a regulatory role. This transcript is present mainly in nucleus, specifically in the chromatin fraction and its silencing results in significant changes in the expression of other *INO80E* and *HIRIP3* transcripts. Thus, based on our experiments, we propose that the INO80E-201 transcript regulates expression of both genes via two alternative but compensatory mechanisms: forming R-loop and forming RNA:RNA duplexes. Both mechanisms might contribute to keeping the chromatin more open and in consequence prevent transcriptional interference. The first 432 nucleotides of INO80E-201 are complementary to *HIRIP3* transcripts and can form a RNA:RNA duplex (Figure 6A). A big part of this region, nucleotides from 51 to 432, is also involved in R-loop formation. Therefore, it is very probable that these two structures have the same function, i.e. the maintenance of a more open chromatin, but are mutually exclusive.

**Figure 6.**
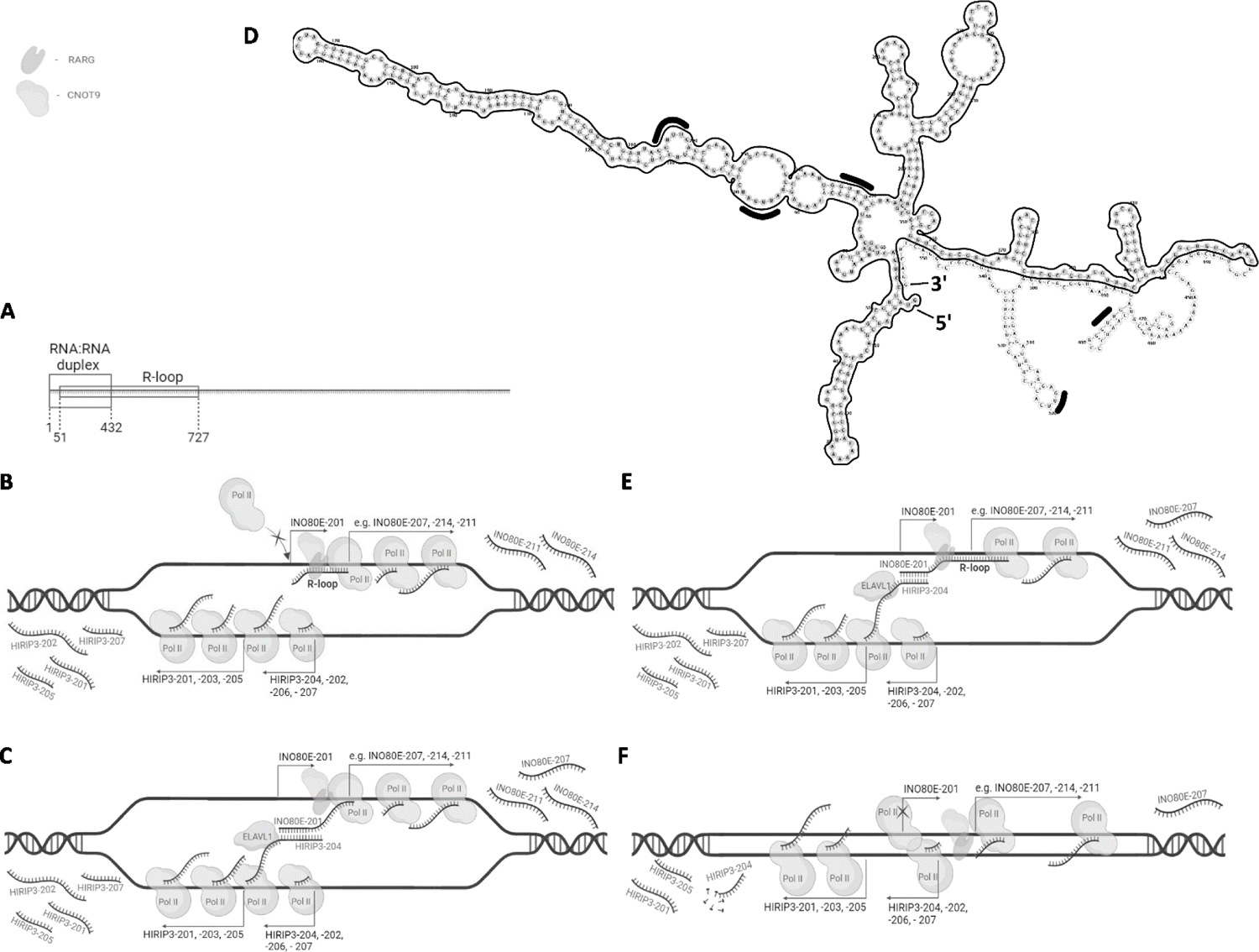
Mechanism of overlapping INO80E and HIRIP3 expression.

Transcription of INO80E-201 requires the binding of a RARG transcription factor that interacts with CNOT9. When an R-loop is formed, transcription of INO80E-201 is blocked, which gives space for polymerases to transcribe *HIRIP3* and non-overlapping transcripts of *INO80E* (Figure 6B). R-loops are not permanent structures and are formed sporadically. Therefore, when an R-loop is not formed or is resolved, the 5’UTR end is available for interaction with the HIRIP3 transcript and form an RNA:RNA duplex. The created duplex stabilizes involved transcripts. Furthermore, the mRNA stabilizing protein ELAVL1 is binds to *HIRIP3* transcripts near the RNA:RNA duplex (Figure 6C). It is also possible that the binding of ELAVL1 prevents the 5’ end of the *HIRIP3* gene from forming secondary structure, which otherwise would block duplex formation (Figure 6D).

Alternatively, both structures could be required at the same time. It is known that R-loops stimulate transcription on the opposite DNA strand (Boque-Sastre et al. 2015; Rondón and Aguilera 2019; Tan-Wong, Dhir, and Proudfoot 2019). Therefore, an R-loop formed by the INO80E-201 transcript may induce a transcription of overlapping *HIRIP3* transcripts. In turn, transcripts of *HIRIP3* form a duplex with INO80E-201, either with its very end (Figure 6E) or with the longer fragment after an R-loop is resolved (Figure 6C). Such a structure involving an R-loop and a dsRNA was already described by the Proudfoot group (Tan-Wong, Dhir, and Proudfoot 2019) as regulating transcription termination in the case of the mouse β-actin gene. However, overexpression of RNAase H1 had no impact on gene expression. This suggests that the forming duplexes might be more crucial and sufficient for keeping chromatin open and for normal genes transcription. Forming R-loops probably plays only a supplementary role and is not required.

Keeping chromatin wide open is essential for undisturbed transcription. This is even more important in the case of 5’ overlapping genes, where transcriptional interference is very likely to occur. A lack of expression of INO80E-201 is equivalent to a lack of RNA:RNA duplexes or R-loops. In consequence, the opening of chromatin is less stable, and polymerase cannot freely either bind to or move in the promoter region (Figure 6F). At the same time however, decreased transcription of overlapping *HIRIP3* isoforms makes downstream located TSS available for polymerase, which results in an increased transcription of HIRIP-201.

## Discussion

Overlapping head-to-head protein-coding genes occur in most organisms, including humans. The implications of such an arrangement of genes are at both the DNA and RNA levels. One possible consequence is that TI may cause a decrease in the level of gene expression, however, there are studies that do not confirm such a pattern (C. H. Chen, Pan, and Lin 2019; Ning et al. 2017; Zhou and Blumberg 2003). Our previous analyses have shown that expression of genes utilizing an TSS which is overlapping at the 5’ end with gene on the opposite DNA strand, is on average higher than when the same genes use only none overlapping TSSs (Rosikiewicz et al. 2021).

In this paper, we present the results of studies on overlapping genes *INO80E* and *HIRIP3*. Both play key roles in the cell and our analysis shows a positive correlation of these two gene expressions. The *INO80E* gene has, according to the available databases (Ensembl, NCBI), one transcript extended at the 5’ end, INO80E-201, which overlaps with the four transcripts of the *HIRIP3* gene (HIRIP3-204, HIRIP3-202, HIRIP3-206, and HIRIP3-207). Our main goal was to determine what role the overlapping transcripts play in regulating the expression of these two genes. The INO80E-201 transcript has an open reading frame and is annotated in databases as encoding the protein. However, our results show that its expression is very low in relation to the other transcripts of the *INO80E* gene (1:766) and it is located in the nucleus. This suggests that the INO80E-201 transcript may have a regulatory function. Transcripts of the *HIRIP3* gene that overlap with *INO80E* are chromatin-associated, although they are also present in other fractions.

Several of our experiments indicated that a decreased expression of INO80E-201 is associated with a decreased expression of remaining *INO80E* transcripts as well as overlapping transcripts of *HIRIP3*. At the same time, the expression level of HIRIP3-201 increased. Changes in the expression of the two remaining *HIRIP3* transcripts, 203 and 205, were not significant. We wanted to check if the introduction of methylation within the promoter sequence of the *INO80E* gene would have similar implications. We introduced *de novo* methylation using the CRISPR/Cas system, consisting of the inactive nuclease dCas9 linked to the DNMT3A protein. As a result of the introduced changes, reduced expression of *HIRIP3* and *INO80E* gene transcripts was observed in the modified cell lines. It was determined that methylated cytosines are located in the binding site for four transcription factors including RARG. It has been described that RARG can activate gene transcription in the presence or in the absence of hormone ligands (Hauksdottir, Farboud, and Privalsky 2003). After a knock-down of *RARG* expression, a similar expression pattern of *INO80E* and *HIRIP3* gene transcripts was observed as after a silencing of the INO80E-201 transcript. Thus, RARG is involved in the regulation of *INO80E* and *HIRIP3* gene expression. Methylation of the *INO80E* gene promoter sequence resulted in a decrease in the expression of the HIRIP3-201 transcript, which is opposite to INO80E-201 silencing. However, this result is not surprising since the methylation process affects the packing of chromatin, leading to a possible silencing of the entire DNA region. It is also possible that the obtained methylations blocked the binding of proteins other than RARG, which affected the expression of HIRIP3-201. It has been proven that cytosine methylation in DNA affects protein binding, including the inhibition of the binding of some transcription factors (Héberlé and Bardet 2019; Yin et al. 2017; Zhu, Wang, and Qian 2016).

We determined that INO80E-201 forms an R-loop and this could positively regulate antisense *HIRIP3* transcripts. This type of mechanism has already been observed and described in the case of other genes (Boque-Sastre et al. 2015; Rondón and Aguilera 2019; Tan-Wong, Dhir, and Proudfoot 2019). R-loop formation at the 5’ end by the INO80E-201 transcript was confirmed by native bisulfite conversion sequencing and supported by RAP-MS results, where proteins responsible for R-loop regulation were identified, including the PARP1 protein (Dou et al. 2020; Lin et al. 2022; Patel et al. 2021; Pérez-Calero et al. 2020; Wolak et al. 2020; Wood et al. 2020). However, RNase H1 overexpression experiments showed that in the lack of an R-loop, expression of all overlapping transcripts increased. This was expected in the case of INO80E-201 since R-loops block transcription elongation but less so expected in the case of overlapping *HIRIP3* transcripts.

We also demonstrated by using RAP-PCR that the INO80E-201 transcript forms an RNA:RNA duplex with overlapping *HIRIP3* transcripts. We propose that the created dsRNA structure acts as a stabilizer of the open loop of DNA, enabling the transcription of the remaining transcripts of both genes. The stabilizing effect of the RNA:RNA duplex was confirmed by transcription termination in HEK293T cells using actinomycin D. The results of this experiment showed that the extended *HIRIP3* gene transcripts degrade faster in INO80E-201 transcript knock-down cells. Forming dsRNA may additionally stabilize overlapping transcripts of the *HIRIP3* gene through the suppression of the mRNA folding at the 5’ end, which in turn may affect the exposure of protein binding sites. Using the available Scan for Motifs database, we found motifs characteristic for the *ELAVL1* protein that we previously identified in RAP-MS. These motifs are located in the 5’UTR of the *HIRIP3* gene, downstream to the sequence involved in the formation of the RNA:RNA duplex. This binding site could be masked by a hairpin formed at the 5’ end of *HIRIP3* transcripts. Although the ELAVL1 protein is most often described in its role of stabilizing transcripts after interaction with the 3’UTR, there are also reports describing a similar function of this protein by interacting with 5’UTR (X. Chen et al. 2019; I. X. Wang et al. 2018). The presented results of the *ELAVL1* gene silencing showed that this protein affects the stability of the transcripts of both genes.

Subsequently, the results of the *HIRIP3* gene expression knock-down showed that the overlapping *HIRIP3* transcripts may also have a stabilizing function for the INO80E-201 transcript, as its expression decreased. However, the expression of the remaining *INO80E* gene transcripts increases upon silencing of the *HIRIP3* gene. These results may indicate that silencing of *HIRIP3* made a promoter region more available for polymerases transcription of none overlapping *INO80E* isoforms. With a significantly lower possibility of transcriptional interference, the presence of an RNA:RNA duplex that maintains the chromatin more openly might not be required for a higher expression. Furthermore, the knock-down of the elongated INO80E-201 transcript resulted in a decreased expression of none overlapping *INO80E* transcripts. This may indicate that in the absence of a RNA:RNA duplex that opens up DNA and stabilizes transcripts, there is not enough space for polymerases that get stacked in the promoter region. Transcription interference, i.e., polymerase pausing in the promoter region, was confirmed using ChIP with an antibody directed to RNA polymerase II. We also noted a diminishment of RNA polymerase II along the *INO80E* gene body and an enrichment in the region of the HIRIP3-201 transcript, for which we observed an increased expression. It is plausible that the lack of a stabilizing duplex facilitates the degradation of overlapping *HIRIP3* transcripts, which makes the more downstream TSS of HIRIP3-201 more accessible for polymerase and would explain an increased expression of this transcript.

As already mentioned, our experiments demonstrated that a decreased expression of INO80E-201 is associated with a decreased expression of the overlapping transcripts of *HIRIP3*. Similar results were obtained by Mahmoudi et al. (Mahmoudi et al. 2009), who studied two protein-coding genes: the *TP53* gene and antisense *WRAP53.* siRNA knockdown of WRAP53 resulted in a significant decrease in *TP53* mRNA. Conversely, overexpression of *WRAP53* increased *TP53* mRNA and p53 protein levels. A blocking of potential *WRAP53/TP53* RNA hybrids also resulted in a reduced *TP53* level. These results strongly suggest that *WRAP53* regulates *TP53* via *WRAP53*/*TP53* RNA interaction.

Altogether, our results strongly suggest that the formation of an RNA:RNA duplex is necessary for stable simultaneous expression of both genes. Lack of this dsRNA structure results in a loss of a stable DNA opening and in consequence transcriptional interference, such as described in the proposed model. These results also explain our previous finding that utilizing an overlapping TSS is associated with a higher expression (Rosikiewicz et al. 2021). They also confirm that transcriptional interference may occur in the case of overlapping genes, but more so in the situation when a non-overlapping TSS are used.

## Materials and methods

### Bioinformatics analysis

*In silico* analyses were carried out using the results of expression estimation studies performed in our laboratory (Szcześniak et al. 2020). The following calculations were performed using 784 ENCODE libraries obtained from normal and cancer tissue/cell line samples samples. Using the Scan for Motifs bioinformatics tool (Biswas and Brown 2014) and RNA-Binding Protein DataBase (RBPDB) (Cook et al. 2011), we confirmed the presence of ELAVL1 protein binding motifs in the 5’UTR sequence of the *HIRIP3* gene. The possibility of R-loops formation within the studied pair of genes, *INO80E* and *HIRIP3*, was checked in the R-loopBase database (Lin et al. 2022). The prediction of DNA binding proteins/transcription factors was made using a database ReMap Atlas of Regulatory Regions at UCSC Genome Browser (Kent et al. 2002) and JASPAR database (Castro-Mondragon et al. 2022). Furthermore, the RNAfold online tool (Kerpedjiev, Hammer, and Hofacker 2015) was used to predict the secondary structure of the overlapping end of the *HIRIP3* gene.

### AMT-crosslinking

HEK293T cells were cultured to the ∼80% confluency (∼25 million cells) on two separate 15-cm plates – one for crosslinked (+4’-aminomethyl trioxsalen (AMT)) and mock-crosslinked (+PBS) conditions. Trypsinized and centrifuged cells were washed once with room temperature PBS, then centrifuged again and resuspened in 4 mL of ice-cold 0.5 mg/ml AMT solution in PBS (+AMT sample) or ice-cold PBS alone (-AMT control sample). Both samples were incubated on ice for fifteen min and were transferred to a pre-chilled 10-cm tissue culture dishes. The crosslinking was conducted on ice, 3-4 cm away from the light source under a long-wave UV bulb (350 nm) in a UV Stratalinker 2400 (Stratagene) for seven minutes with mixing breaks every two minutes. After this procedure, the cells were centrifuged 330× g for four minutes and the cell pellets were used directly in the standard RNA isolation procedure.

### RNA isolation, RT-PCR and PCR

Total, cytoplasmic, nucleoplasmic, total nuclear, and chromatin RNA was extracted using TRI Reagent® (Molecular Research Center) with a standard protocol. RNA yield was measured with a DeNovix DS-11 Series spectrophotometer (DeNovix). RNA was reverse transcribed with the RevertAid First Strand cDNA Synthesis Kit (ThermoFisher Scientific) according to the manufacturer’s protocol. Standard PCRs were performed using StartWarm HS-PCR Mix (A&A Biotechnology) or EconoTaq PLUS2X Master Mix (Lucigen). PCR products were analyzed with 1.5% agarose gel containing GelRed (Biotium) in a 1X TAE buffer and subsequently captured the gel images using G:Box EF2 (Syngene) with GeneSys image analysis software (Syngene).

### RNA Antisense Purification (RAP)

Biotinylated probes and RAP were performed according to the described protocol (Engreitz, Lander, and Guttman 2015). In brief, samples were incubated for 2 h with denaturized 15 pmol of biotinylated ssDNA probes. Next, samples were incubated for thirty minutes with subsequently prepared Dynabeads™ MyOne™ Streptavidin C1 magnetic beads (Invitrogen™, ThermoFisher Scientific). Magnetic beads with captured probe:target transcript complexes were purified by washing: three times with Low Stringency Wash Buffer (1× SSPE, 0.1% SDS, 1% NP-40, 4 M urea), three times with High Stringency Wash Buffer (0.1× SSPE, 0.1% SDS, 1% NP-40, 4 M urea), and two times with RNase H Elution Buffer (50 mM Tris-HCl pH 7.5, 75 mM NaCl, 3 mM MgCl2, 0.125% N-lauroylsarcosine, 0.025% sodium deoxycholate, 2.5 mM TCEP). To remove the ssDNA probe bond with RNA, samples were incubated with RNase H (ThermoFisher Scientific) at 37°C for thirty minutes. Eluted RNA was cleaned with the standard TRI Reagent® (Molecular Research Center) extraction.

### RAP-MS

RAP-MS was performed according to the described protocol (Engreitz, Lander, and Guttman 2015). Concisely, cells were UV crosslinked at the 254 nm wavelength with 0.8 J/cm^2^. Then, the cell pellet was resuspended in 1 ml of Cell Lysis Buffer - Nuclear I (10 mM HEPES pH 7.4, 20 mM KCl, 1.5 mM MgCl2, 0.5 mM EDTA, 1 mM TCEP, 0.5 mM PMSF). The MS analysis was performed in the Mass Spectrometry Laboratory, IBB PAS, Warsaw.

### Cell culture

The experiments were performed on HEK239T cells (CRL-1573™, ATCC). Cells were maintained under standard conditions at 37°C in a humidified incubator with 5% CO2 in DMEM (Genos) supplemented with 10% fetal bovine serum – FBS (Genos) and 1 % Penicillin-Streptomycin (10,000 U/mL) (Gibco™, ThermoFisher Scientific). Cells were trypsinized using Trypsin (0.25%) (Gibco™, ThermoFisher Scientific).

### RNA knock-down protocol

The silencing of the selected transcripts was performed using Antisense LNA GapmeRs (Qiagen) or Silencer Select siRNAs (ThermoFisher). For Antisense LNA GapmeRs (Qiagen) silencing the 30 nM of GapmeRs were used and for Silencer Select siRNAs silencing the 75 nM of siRNA. Cells after transfection were cultured for 72 h, then were collected and used for RNA or protein extraction.

### Cell transfection

Genetic material was delivered to cells using Lipofectamine™ 3000 Transfection Reagent (Invitrogen™, ThermoFisher Scientific). Prior transfection cells were cultured to the ∼70% confluency on 12-well dishes. Lipofectamine™ 3000 in an amount of 3 µl was diluted in 50 µl of Opti-MEM™ I Reduced Serum Medium, no phenol red) (Gibco™, ThermoFisher Scientific) per one well. Transfection material was diluted in 50 µl of Opti-MEM™ I Reduced Serum Medium, no phenol red) (Gibco™, ThermoFisher Scientific) and addition of 2 µl of P3000™ Reagent per one well. Then, diluted Lipofectamine™ 3000 and a transfection material were mixed and were incubated for fifteen minutes at room temperature. After incubation the lipid complexes were added to the cells carefully.

### Subcellular fractionation

Isolation of RNA from cytoplasmic, total nuclear, nucleoplasmic, and chromatin fractions were performed from cells cultured to the ∼90% confluency on 10 cm plates according to the described protocol (Gagnon et al. 2014). In brief, collected cells were resuspended in 380 μl of ice-cold Hypotonic lysis buffer (10 mM Tris pH 7.5, 10 mM NaCl, 3 mM MgCl2, 0.3% NP-40 and 10% glycerol) supplemented with 100 U of RiboLock RNase Inhibitor (ThermoFisher Scientific) and were incubated on ice for ten minutes. After centrifugation (200 x g at 4°C for 2 min), supernatants were mixed with 1 ml of RNA precipitation solution (5% 3 M sodium acetate (pH 5.5) in 99,6% ethanol) and stored in −20°C for at least 1 h. Pellets were washed three times with 1 ml of ice-cold Hypotonic lysis buffer. Total nuclear RNA samples were suspended in TRI Reagent® (Molecular Research Center) for direct RNA extraction. Pellets for the isolation of nucleoplasmic and chromatin fractions were suspended in 380 μl of Modified Wuarin-Schibler buffer (10 mM Tris-HCl pH 7.0, 4 mM EDTA, 0.3 M NaCl, 1 M urea, and 1% NP-40) supplemented with 100 U of RiboLock RNase Inhibitor (ThermoFisher Scientific) and incubated on ice for ten minutes. After centrifugation (1,000 x g at 4°C for three minutes), supernatants were mixed with 1 ml of RNA precipitation solution and stored as described above. Pellets were washed three times with 1 ml of ice-cold Modified Wuarin-Schibler buffer. Chromatin samples were suspended in TRI Reagent® (Molecular Research Center) for direct RNA extraction. Samples incubated in RNA precipitation solution were centrifuged at 18,000 x g at 4°C for fifteen minutes and washed in ice-cold 70% ethanol. Semi-dry pellets were suspended in TRI Reagent® (Molecular Research Center) for direct RNA extraction.

### Quantitative PCR

Extracted total, cytoplasmic, total nuclear, nucleoplasmic, and chromatin RNA was used for cDNA synthesis using the RevertAid First Strand cDNA Synthesis Kit (Thermo Scientific) according to the manufacturer’s procedure. Quantitative PCR was performed using PowerUp™ SYBR™ Green Master Mix (Applied Biosystems™, ThermoFisher Scientific) on a QuantStudio™ 7 Flex Real-Time PCR System platform (ThermoFisher Scientific). All experiments were carried out in triplicate technical repeats for three biological replicates. For the expression normalization, as an endogenous control, the expression of GAPDH was measured. Then, the data were analyzed using the 2−ΔΔCt method. The statistical significance of the results was performed in GraphPad Prism 9 (GraphPad Software).

### Western Blotting

Cellular extracts for western blotting were prepared from the ∼90% confluency 10 cm plates. After decanting the medium, cells were washed twice with ice-cold PBS and were covered with the ice-cold RIP buffer (150 mM KCl, 25 mM TRIS pH 8, 5 mM EDTA, 0.5 mM DTT, 0.5% Igepal, cOmplete™, Mini Protease Inhibitor Cocktail (Roche, Merck). Then cells were scraped and incubated for 30 minutes at 4°C. Sonication of the samples was carried out for ten minutes (30s ON – 30s OFF intervals) on Bioruptor® (Diagenode Diagnostics). The estimation of the protein amount using Pierce™ BCA Protein Assay Kit (ThermoFisher Scientific) were performed on supernatant after samples centrifugation at 13000 rpm for twenty min at 4 °C. The 30 µg of protein was separated in a 5% stacking gel at 125V and 10% separation gel at 160 V. Then, gels were rinsed with Towbin buffer (25 mM Tris, 192 mM glycine, 20% methanol) and transferred in BioRad Trans-Blot Turbo (Bio-Rad) for thirty minutes at 15 V. Blocked and rinsed membranes were incubated overnight at 4°C with INO80E Polyclonal Antibody (Invitrogen, ThermoFisher Scientific) at concentration 0.04 µg/mL. The membrane was then incubated with IgG (HRP) secondary antibody (ab205718) (Abcam). Membranes were visualized using SuperSignal™ West Femto Maximum Sensitivity Substrate (Thermo Scientific).

### CRISPR/Cas9 methylation

*De novo* methylation was obtained with the use of pdCas9-DNMT3A-PuroR_v2, which was a gift from Vlatka Zoldoš (Addgene plasmid #74407; http://n2t.net/addgene:74407; RRID: Addgene_74407). sgRNA oligonucleotides were designed on the Benchling platform and ordered from Merck. Oligonucleotides were annealed and phosphorylated with 5U of T4 Polynucleotide Kinase (New England Biolabs). Methylation constructs were obtained with the Golden Gate assembly cloning strategy based on *Bbs*I restriction enzyme. Obtained constructs were transformed into DH5α *E.coli* cells. Positive clones, after colony PCR check, were cultured for isolation with Plasmid Mini Kit (A&A Biotechnology). Properly cloned constructs were verified using Sanger sequencing.

### Bisulfite conversion

Bisulfite conversion was done with the EpiJET Bisulfite Conversion Kit (Thermo Scientific) according to the manufacturer protocol. In brief, samples with 500 ng of DNA were mixed with 120 µl freshly prepared Modification Reagent, then incubated in 98°C for ten minutes and 60°C for 150 minutes. Converted DNA samples were purified and used for PCR reactions.

### RNase H overexpression

The pEGFP-RNASEH1 was a gift from Andrew Jackson & Martin Reijns (Addgene plasmid #108699; http://n2t.net/addgene:108699; RRID:Addgene_108699). Plasmids were introduced to the HEK293T cells with the Lipofectamine™ 3000 Transfection Reagent (see: Cell transfection). Transfection success was checked after 48h using a ZOE Fluorescent Cell Imager (Bio-Rad). Fluorescence-activated cell sorting (FACS) was then performed to separate GFP-positive cells that had taken up the pEGFP-RNASEH1 plasmid.

### Chromatin Immunoprecipitation

Proteins were cross-linked to DNA with 1% formadehyde. After ten minutes of incubation, cells were quenched with 125 mM glycine. Then, cells were scraped and incubated for five minutes at 4°C with two lysis buffers (L1: 50 mM TRIS pH 8, 1 mM EDTA pH 8, 0.1% NP40, 10% glycerol, Protease Inhibitors; L2: 1% SDS, 50 mM TRIS pH 8, 10 mM EDTA pH 8, Protease Inhibitors). Sonication of the samples was carried out for ten minutes (30s ON – 30s OFF intervals) on Bioruptor® (Diagenode Diagnostics). The chromatins were checked with gel electrophoresis. Chromatin immunoprecipitation was performed with the use of DynaBeads protein A/G (Thermo Scientific), histone H3 antibody - (ab1791) Anti-Histone H3 antibody (Abcam), and RNA polymerase II antibody - (ab10338) Anti-RPB2 antibody (Abcam).

### R-loop Native Bisulfite Sequencing – chromatin preparation

This experiment was based on a protocol published by Boque-Sastre and colleques (Boque-Sastre, Soler, and Guil 2017). In brief, cells at ∼80% confluency from 6-well plate were scraped and centrifuged. Cell pellets were incubated in 500 µl of lysis buffer (10 mM TRIS pH 8, 1 mM EDTA, 0.5% SDS with 55 µg Proteinase K) at 37°C overnight. Cell lysates was mixed with 250 µl 5 M NaCl and centrifuged at maximum speed for fifteen minutes. Precipitated chromatin was used for bisulfite conversion.

### pGEM-T cloning of PCR products

The pGEM®-T Easy Vector System (Promega) was used according to the manufacturer protocol. In brief, the standard ligation reaction mixes were incubated overnight at 4°C. Next, the obtained vectors were mixed with insert in 1:3 ratio and transformed into DH5α *E.coli* cells. Selection for transformants was conducted on LB/ampicillin/IPTG/X-Gal plates. Positive, white-color clones were cultured for isolation with Plasmid Mini Kit (A&A Biotechnology). Vectors were verified using Sanger sequencing.

### Statistical analysis

Expression values were examined using GraphPad Prism 9 (GraphPad, San Diego, CA, USA, www.graphpad.com) and R (R Core Team 2013) with the “ggsignif” (Ahlmann-Eltze and Patil 2023), “ggplot2” (Wickham 2009), “ggh4x” (Brand 2022), “smplot2” (Seung 2023), “tidyverse” (Wickham et al. 2019), and “ggpubr” (Kassambara 2023) packages. Firstly, it was determined whether data is normally distributed with the Shapiro-Wilk normality test. Then, the Mann-Whitney U test was done in order to compare differences between expression levels of different overlapping genes and transcripts. The relation between *INO80E* and *HIRIP3* expression levels was studied using the Spearman correlation test. In all analyses, p < 0.05 was used as statistically significant. Statistical analysis of relative expressions of tested transcripts after qPCR was performed in GraphPad Prism 9 (GraphPad, San Diego, CA, USA, www.graphpad.com). The Brown-Forsythe and Welch ANOVA test with Dunnett’s T3 multiple comparisons tests were used.

