## Supplementary Materials for "5’ extended protein-coding *INO80E* transcript regulates expression of two head-to-head overlapping genes: *INO80E* and *HIRIP3*"

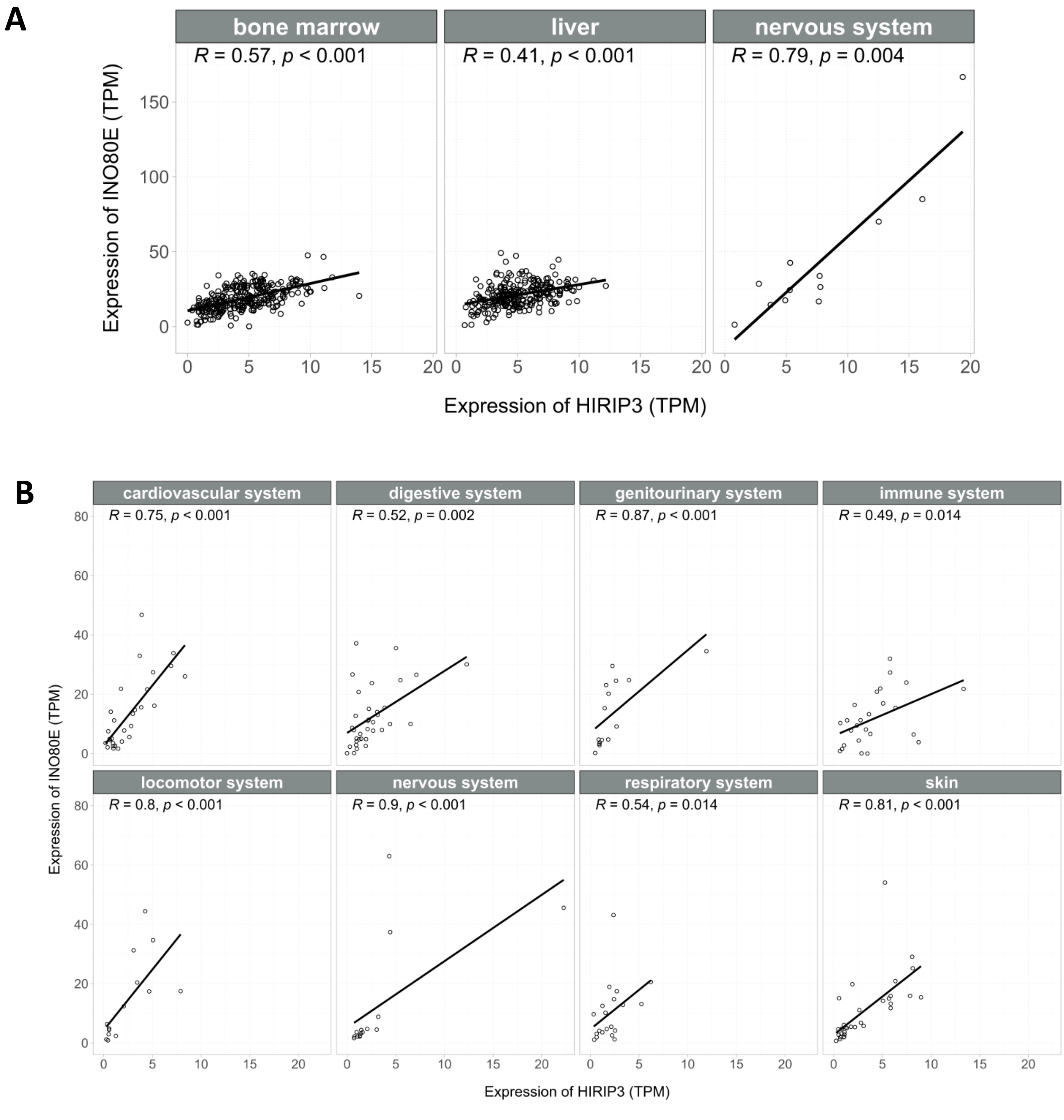

**Figure S1.** Correlation between *INO80E* gene and *HIRIP3* gene expression (TPM): A - in bone marrow, liver and brain tumor cancer cells; C - in non-cancer cardiovascular, digestive, genitourinary, immune, locomotor, nervous, respiratory system and skin cells.

**A**

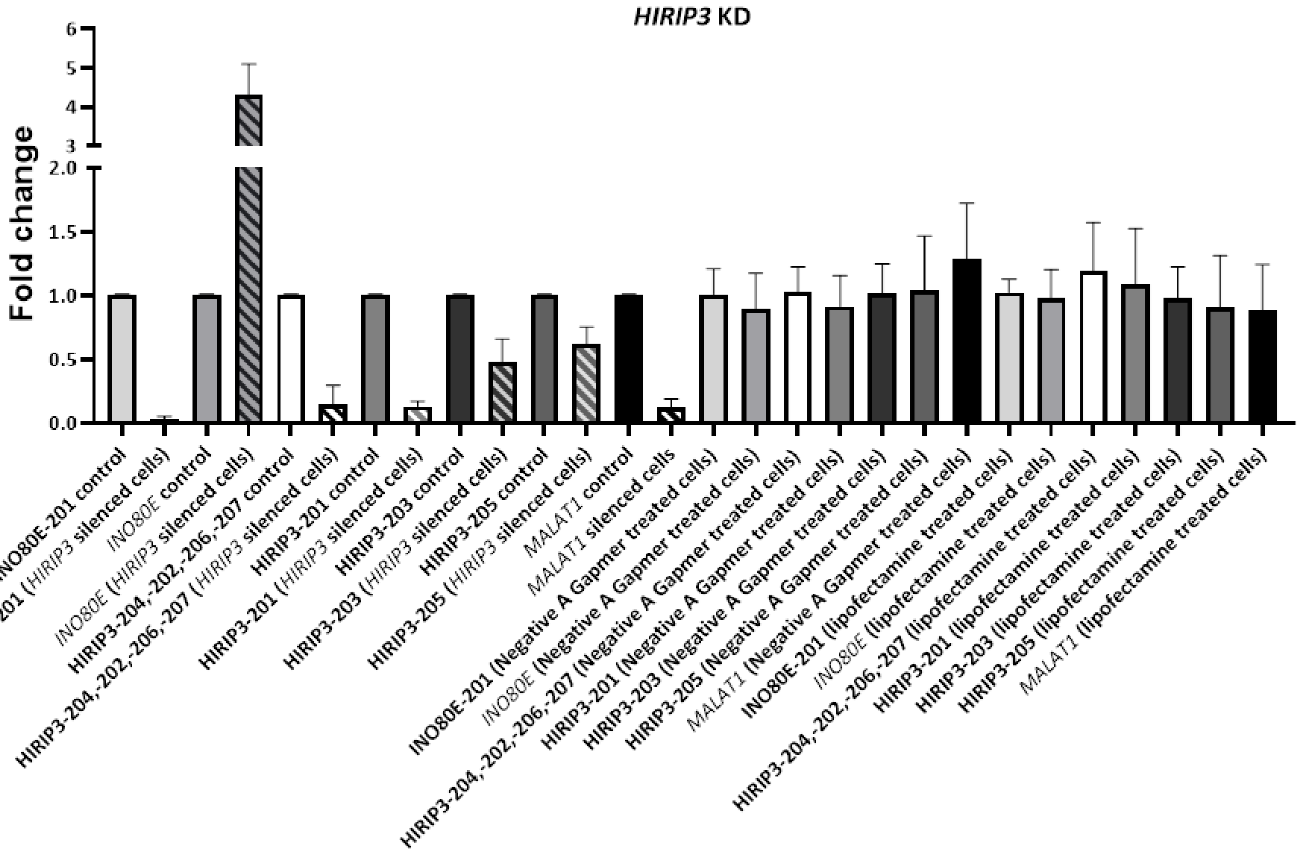

| **Brown-Forsythe and Welch ANOVA test; Dunnett's T3 multiple comparisons test** | **Summary** | **Adjusted P Value** |
|---|---|---|
| INO80E-201 control vs. INO80E-201 (HIRIP3 silenced cells) | **** | <0,0001 |
| INO80E-201 control vs. INO80E-201 (Negative A Gapmer treated cells) | ns | >0,9999 |
| INO80E-201 control vs. INO80E-201 (lipofectamine treated cells) | ns | >0,9999 |
| INO80E control vs. INO80E (HIRIP3 silenced cells) | *** | 0,0002 |
| INO80E control vs. INO80E (Negative A Gapmer treated cells) | ns | >0,9999 |
| INO80E control vs. INO80E (lipofectamine treated cells) | ns | >0,9999 |
| HIRIP3-204,-202,-206,-207 control vs. HIRIP3-204,-202,-206,-207 (HIRIP3 silenced cells) | **** | <0,0001 |
| HIRIP3-204,-202,-206,-207 control vs. HIRIP3-204,-202,-206,-207 (Negative A Gapmer treated cells) | ns | >0,9999 |
| HIRIP3-204,-202,-206,-207 control vs. HIRIP3-204,-202,-206,-207 (lipofectamine treated cells) | ns | 0,9994 |
| HIRIP3-201 control vs. HIRIP3-201 (HIRIP3 silenced cells) | **** | <0,0001 |
| HIRIP3-201 control vs. HIRIP3-201 (Negative A Gapmer treated cells) | ns | >0,9999 |
| HIRIP3-201 control vs. HIRIP3-201 (lipofectamine treated cells) | ns | >0,9999 |
| HIRIP3-203 control vs. HIRIP3-203 (HIRIP3 silenced cells) | ** | 0,0028 |
| HIRIP3-203 control vs. HIRIP3-203 (Negative A Gapmer treated cells) | ns | >0,9999 |
| HIRIP3-203 control vs. HIRIP3-203 (lipofectamine treated cells) | ns | >0,9999 |
| HIRIP3-205 control vs. HIRIP3-205 (HIRIP3 silenced cells) | ** | 0,0029 |
| HIRIP3-205 control vs. HIRIP3-205 (Negative A Gapmer treated cells) | ns | >0,9999 |
| HIRIP3-205 control vs. HIRIP3-205 (lipofectamine treated cells) | ns | >0,9999 |
| MALAT1 control vs. MALAT1 silenced cells | **** | <0,0001 |
| MALAT1 control vs. MALAT1 (Negative A Gapmer treated cells) | ns | >0,9999 |
| MALAT1 control vs. MALAT1 (lipofectamine treated cells) | ns | >0,9999 |

**B**

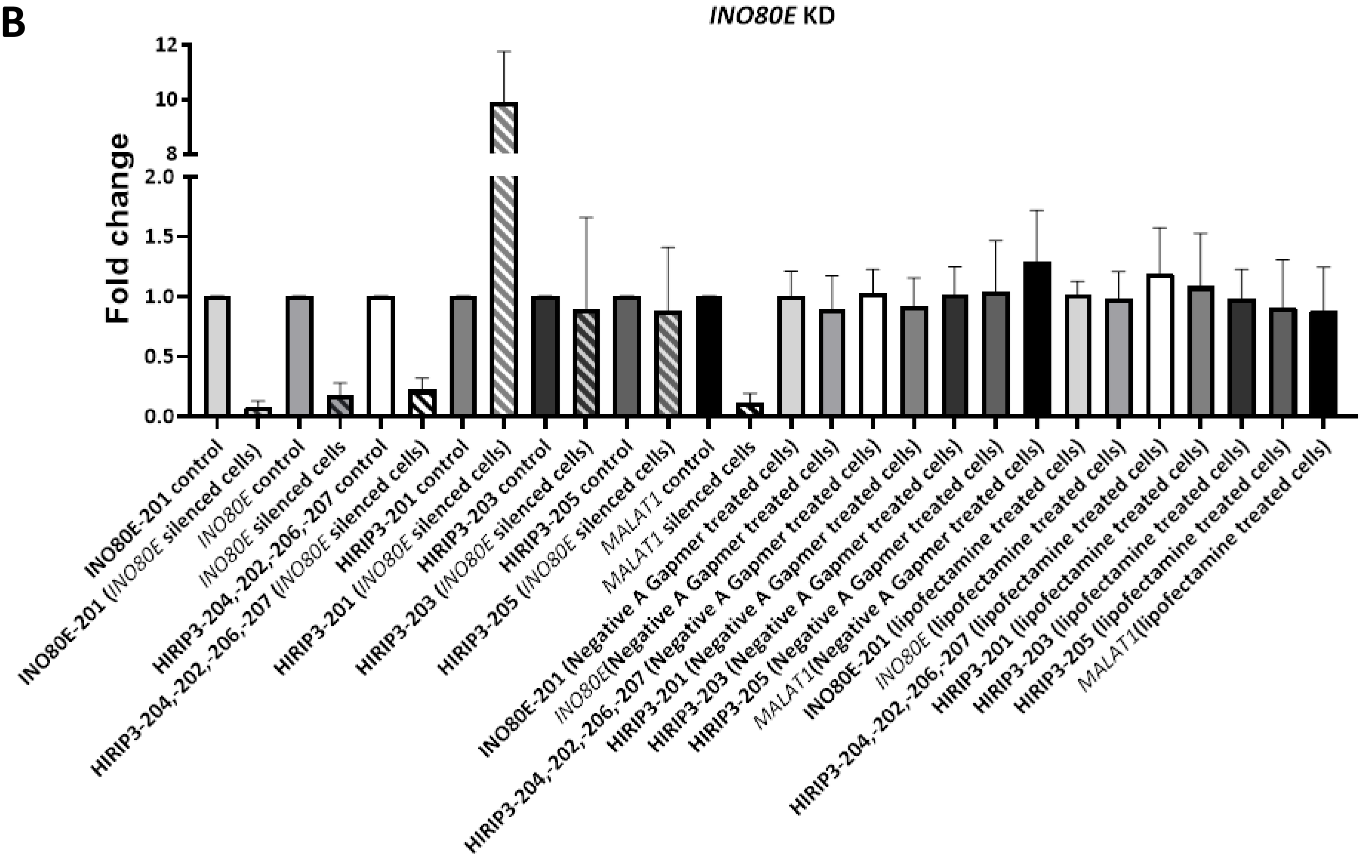

| Brown-Forsythe and Welch ANOVA test; Dunnett's T3 multiple comparisons test | Summary | Adjusted P Value |
|---|---|---|
| INO80E-201 control vs. INO80E-201 (INO80E silenced cells) | **** | <0,0001 |
| INO80E-201 control vs. INO80E-201 (Negative A Gapmer treated cells) | ns | >0,9999 |
| INO80E-201 control vs. INO80E-201 (lipofectamine treated cells) | ns | >0,9999 |
| INO80E control vs. INO80E silenced cells | **** | <0,0001 |
| INO80E control vs. INO80E (Negative A Gapmer treated cells) | ns | >0,9999 |
| INO80E control vs. INO80E (lipofectamine treated cells) | ns | >0,9999 |
| HIRIP3-204,-202,-206,-207 control vs. HIRIP3-204,-202,-206,-207 (INO80E silenced cells) | **** | <0,0001 |
| HIRIP3-204,-202,-206,-207 control vs. HIRIP3-204,-202,-206,-207 (Negative A Gapmer treated cells) | ns | >0,9999 |
| HIRIP3-204,-202,-206,-207 control vs. HIRIP3-204,-202,-206,-207 (lipofectamine treated cells) | ns | 0,9994 |
| HIRIP3-201 control vs. HIRIP3-201 (INO80E silenced cells) | **** | <0,0001 |
| HIRIP3-201 control vs. HIRIP3-201 (Negative A Gapmer treated cells) | ns | >0,9999 |
| HIRIP3-201 control vs. HIRIP3-201 (lipofectamine treated cells) | ns | >0,9999 |
| HIRIP3-203 control vs. HIRIP3-203 (INO80E silenced cells) | ns | >0,9999 |
| HIRIP3-203 control vs. HIRIP3-203 (Negative A Gapmer treated cells) | ns | >0,9999 |
| HIRIP3-203 control vs. HIRIP3-203 (lipofectamine treated cells) | ns | >0,9999 |
| HIRIP3-205 control vs. HIRIP3-205 (INO80E silenced cells) | ns | >0,9999 |
| HIRIP3-205 control vs. HIRIP3-205 (Negative A Gapmer treated cells) | ns | >0,9999 |
| HIRIP3-205 control vs. HIRIP3-205 (lipofectamine treated cells) | ns | >0,9999 |
| MALAT1 control vs. MALAT1 silenced cells | **** | <0,0001 |
| MALAT1 control vs. MALAT1 (Negative A Gapmer treated cells) | ns | >0,9999 |
| MALAT1 control vs. MALAT1 (lipofectamine treated cells) | ns | >0,9999 |

**C**

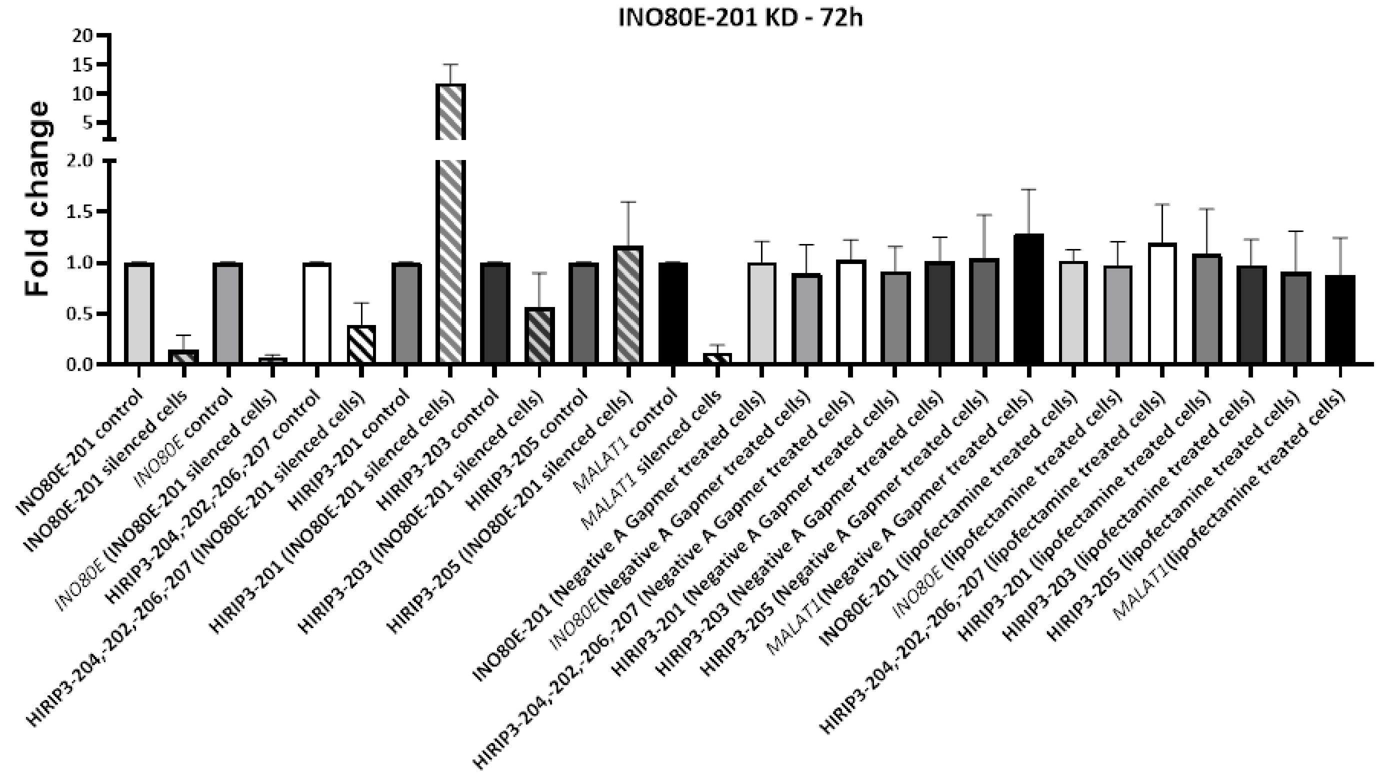

| Brown-Forsythe and Welch ANOVA test; Dunnett's T3 multiple comparisons test | Summary | Adjusted P Value |
|---|---|---|
| INO80E-201 control vs. INO80E-201 silenced cells | **** | <0,0001 |
| INO80E-201 control vs. INO80E-201 (Negative A Gapmer treated cells) | ns | >0,9999 |
| INO80E-201 control vs. INO80E-201 (lipofectamine treated cells) | ns | >0,9999 |
| INO80E control vs. INO80E (INO80E-201 silenced cells) | **** | <0,0001 |
| INO80E control vs. INO80E (Negative A Gapmer treated cells) | ns | >0,9999 |
| INO80E control vs. INO80E (lipofectamine treated cells) | ns | >0,9999 |
| HIRIP3-204,-202,-206,-207 control vs. HIRIP3-204,-202,-206,-207 (INO80E-201 silenced cells) | ** | 0,0039 |
| HIRIP3-204,-202,-206,-207 control vs. HIRIP3-204,-202,-206,-207 (Negative A Gapmer treated cells) | ns | >0,9999 |
| HIRIP3-204,-202,-206,-207 control vs. HIRIP3-204,-202,-206,-207 (lipofectamine treated cells) | ns | 0,9994 |
| HIRIP3-201 control vs. HIRIP3-201 (INO80E-201 silenced cells) | *** | 0,0009 |
| HIRIP3-201 control vs. HIRIP3-201 (Negative A Gapmer treated cells) | ns | >0,9999 |
| HIRIP3-201 control vs. HIRIP3-201 (lipofectamine treated cells) | ns | >0,9999 |
| HIRIP3-203 control vs. HIRIP3-203 (INO80E-201 silenced cells) | ns | 0,2874 |
| HIRIP3-203 control vs. HIRIP3-203 (Negative A Gapmer treated cells) | ns | >0,9999 |
| HIRIP3-203 control vs. HIRIP3-203 (lipofectamine treated cells) | ns | >0,9999 |
| HIRIP3-205 control vs. HIRIP3-205 (INO80E-201 silenced cells) | ns | >0,9999 |
| HIRIP3-205 control vs. HIRIP3-205 (Negative A Gapmer treated cells) | ns | >0,9999 |
| HIRIP3-205 control vs. HIRIP3-205 (lipofectamine treated cells) | ns | >0,9999 |
| MALAT1 control vs. MALAT1 silenced cells | **** | <0,0001 |
| MALAT1 control vs. MALAT1 (Negative A Gapmer treated cells) | ns | >0,9999 |
| MALAT1 control vs. MALAT1 (lipofectamine treated cells) | ns | >0,9999 |

**Figure S2.** Fold change of *HIRIP3* (**A**) and *INO80E* (**B**) transcripts expression involved in the creation of RNA duplex after silencing one gene in a pair at a time and INO80E-201 (**C**). The *MALAT1* silencing is a positive control, Negative A GapmeR silencing is a negative control, lipofectamine treated cells is a control of transfection (ns – not significant; ** - p<0.01; *** - p<0.001; **** - p<0.0001).

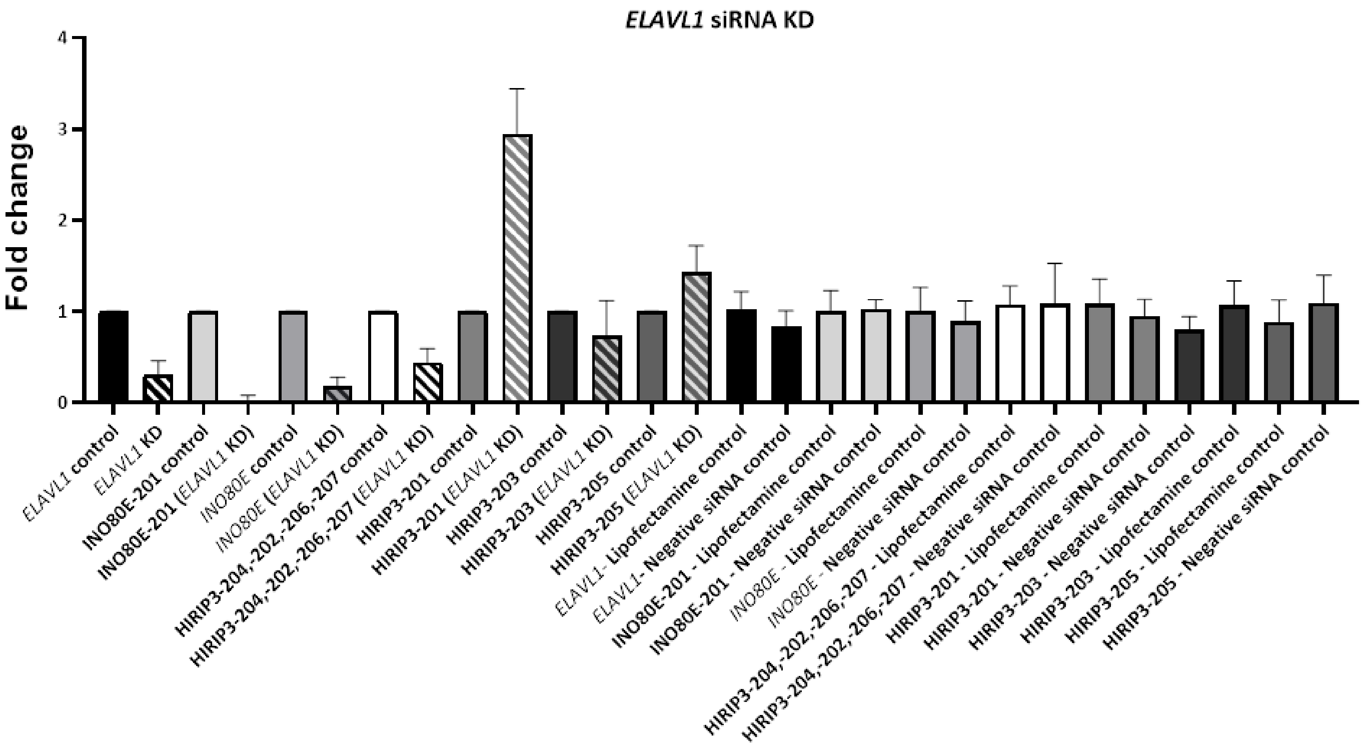

| Dunnett's T3 multiple comparisons test | Summary | Adjusted P Value |
|---|---|---|
| *ELAVL1*control vs. *ELAVL1*KD | *** | 0,0001 |
| *ELAVL1*control vs. *ELAVL1*- Lipofectamine control | ns | >0,9999 |
| *ELAVL1*control vs. *ELAVL1*- Negative siRNA control | ns | 0,7067 |
| INO80E-201 control vs. INO80E-201 (*ELAVL1*KD) | **** | <0,0001 |
| INO80E-201 control vs. INO80E-201 - Lipofectamine control | ns | >0,9999 |
| INO80E-201 control vs. INO80E-201 - Negative siRNA control | ns | >0,9999 |
| *INO80E* control vs. *INO80E* (*ELAVL1*KD) | **** | <0,0001 |
| *INO80E* control vs. *INO80E* - Lipofectamine control | ns | >0,9999 |
| *INO80E* control vs. *INO80E* - Negative siRNA control | ns | 0,9999 |
| HIRIP3-204,-202,-206,-207 control vs. HIRIP3-204,-202,-206,-207 (*ELAVL1* KD) | *** | 0,0005 |
| HIRIP3-204,-202,-206,-207 control vs. HIRIP3-204,-202,-206,-207 - Lipofectamine control | ns | >0,9999 |
| HIRIP3-204,-202,-206,-207 control vs. HIRIP3-204,-202,-206,-207 - Negative siRNA control | ns | >0,9999 |
| HIRIP3-201 control vs. HIRIP3-201 (*ELAVL1*KD) | *** | 0,0003 |
| HIRIP3-201 control vs. HIRIP3-201 - Lipofectamine control | ns | >0,9999 |
| HIRIP3-201 control vs. HIRIP3-201 - Negative siRNA control | ns | >0,9999 |
| HIRIP3-203 control vs. HIRIP3-203 (*ELAVL1*KD) | ns | 0,9686 |
| HIRIP3-203 control vs. HIRIP3-203 - Lipofectamine control | ns | >0,9999 |
| HIRIP3-203 control vs. HIRIP3-203 - Negative siRNA control | ns | 0,1944 |
| HIRIP3-205 control vs. HIRIP3-205 (*ELAVL1*KD) | ns | 0,9686 |
| HIRIP3-205 control vs. HIRIP3-205 - Lipofectamine control | ns | >0,9999 |
| HIRIP3-205 control vs. HIRIP3-205 - Negative siRNA control | ns | 0,9732 |

**Figure S3**. *ELAVL1* gene knock-down effect on the *INO80E* and *HIRIP3* genes. The negative siRNA control silencing is a negative control, lipofectamine treated cells is a control of transfection (ns – not significant; *** - $p<0.001$; **** - $p<0.0001$).

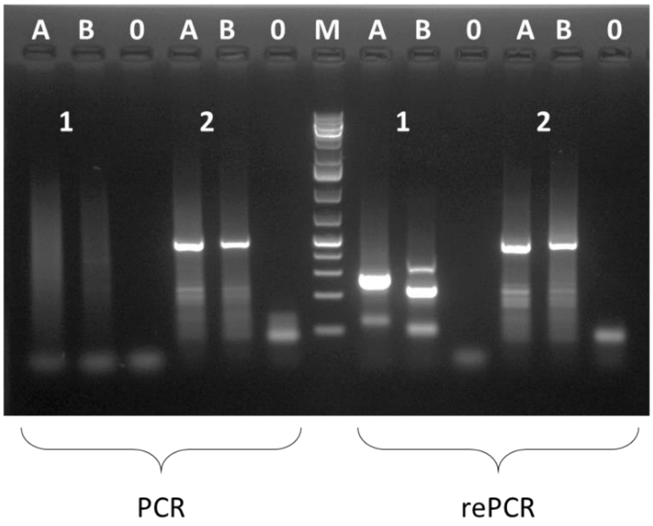

**Figure S4.** R-loop formation within the *INO80E* gene. Results from native bisulfite PCR and rePCR. 1 – primers amplifying the R-loop region and upstream; 2 - control primers amplifying the region within the R-loop; A – control WT HEK293 cells; B – INO80E-201 silenced HEK293 cells; 0 - PCR purity control

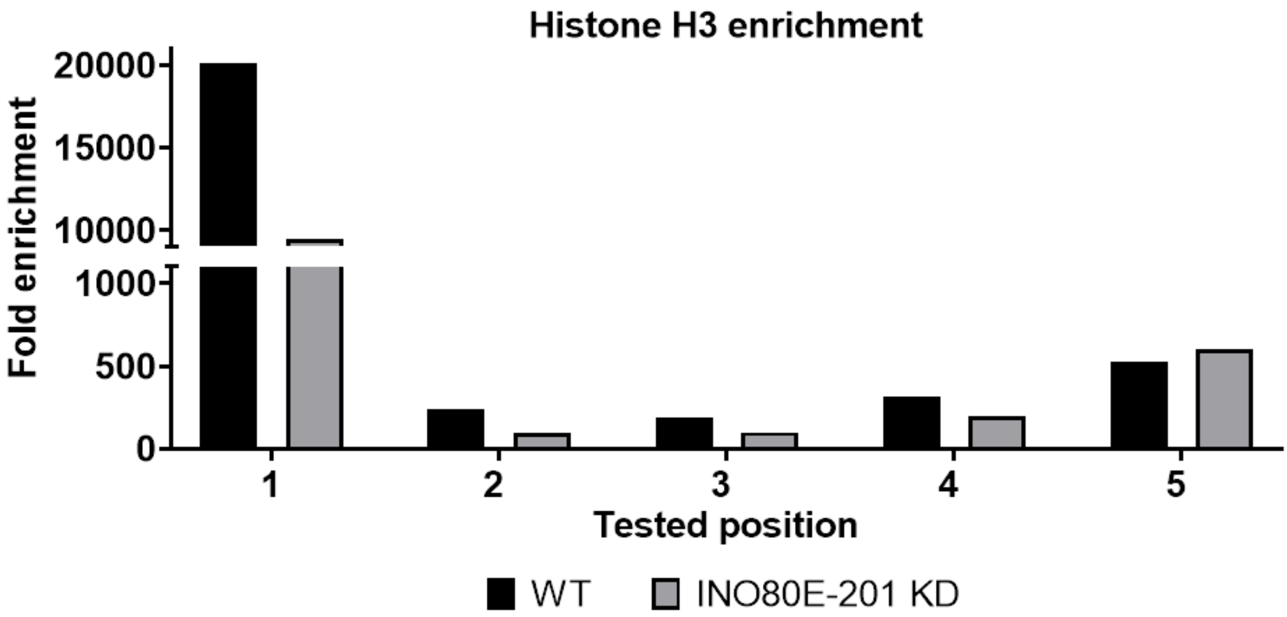

**Figure S5.** Chromatin immunoprecipitation with Histone H3 antibody. The numbers indicates the location of the RNA Polymerase II binding detection primers.

**A**

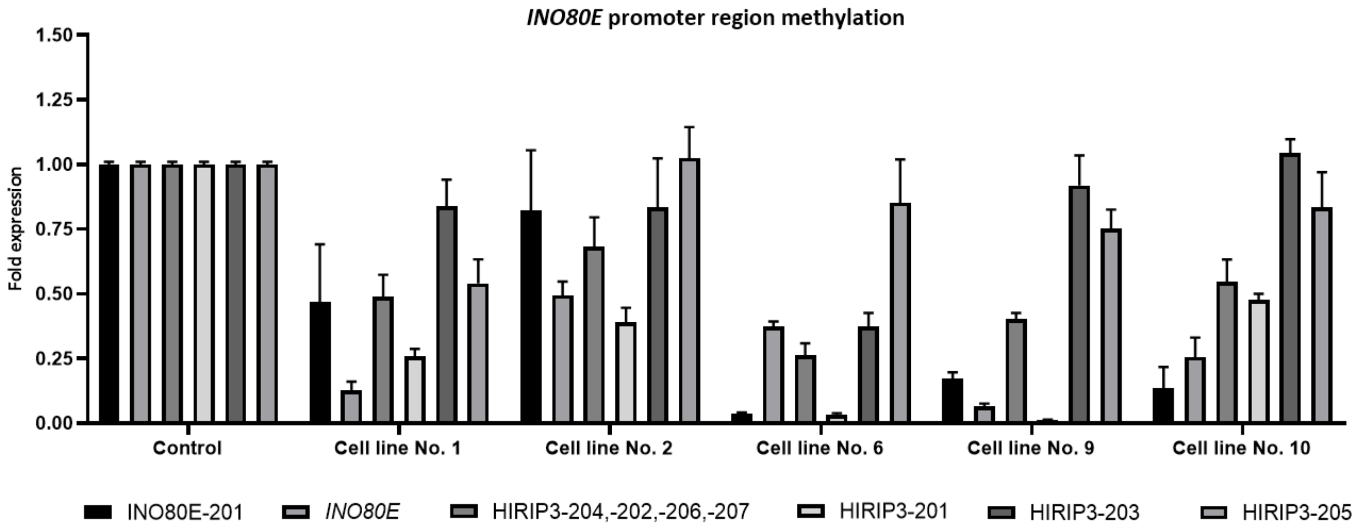

| Tukey's multiple comparisons test | | | |
|---|---|---|---|
| INO80E-201 | | HIRIP3-201 | |
| Control vs. Cell line No. 1 | ns | Control vs. Cell line No. 1 | **** |
| Control vs. Cell line No. 2 | ns | Control vs. Cell line No. 2 | ** |
| Control vs. Cell line No. 6 | **** | Control vs. Cell line No. 6 | **** |
| Control vs. Cell line No. 9 | **** | Control vs. Cell line No. 9 | **** |
| Control vs. Cell line No. 10 | ** | Control vs. Cell line No. 10 | *** |
| *INO80E* | | HIRIP3-203 | |
| Control vs. Cell line No. 1 | **** | Control vs. Cell line No. 1 | ns |
| Control vs. Cell line No. 2 | * | Control vs. Cell line No. 2 | ns |
| Control vs. Cell line No. 6 | **** | Control vs. Cell line No. 6 | ** |
| Control vs. Cell line No. 9 | **** | Control vs. Cell line No. 9 | ns |
| Control vs. Cell line No. 10 | * | Control vs. Cell line No. 10 | ns |
| HIRIP3-204,-202,-206,-207 | | HIRIP3-205 | |
| Control vs. Cell line No. 1 | * | Control vs. Cell line No. 1 | * |
| Control vs. Cell line No. 2 | ns | Control vs. Cell line No. 2 | ns |
| Control vs. Cell line No. 6 | ** | Control vs. Cell line No. 6 | ns |
| Control vs. Cell line No. 9 | **** | Control vs. Cell line No. 9 | ns |
| Control vs. Cell line No. 10 | * | Control vs. Cell line No. 10 | ns |

**B**

| Tested cell line number | Number of identified methylated cytosines |
|---|---|
| 1 | +4 |
| 2 | +16 |
| **6** | +9 |
| **9** | +15 |
| 10 | +9 |

**Figure S6.** *INO80E* promoter region methylation. A: The fold expression of *INO80E* and *HIRIP3* genes transcripts with the table of the statistical significance of the results. B: Number of identified methylated cytosines in tested cell lines (ns – not significant, *- p<0.05; **- p<0.01; **** - p<0.0001).

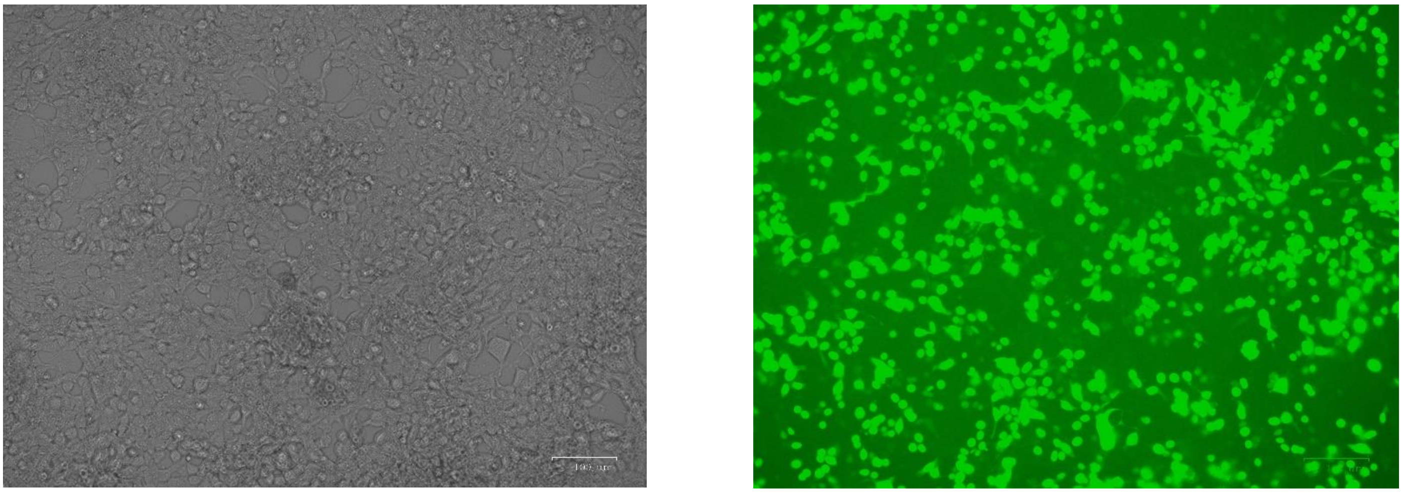

**Figure S7.** RNase H1 overexpression images from ZOE Fluorescent Cell Imager of the transfected cells 48 hours after transfection (left – brightfield, right - green channel using a blue LED).

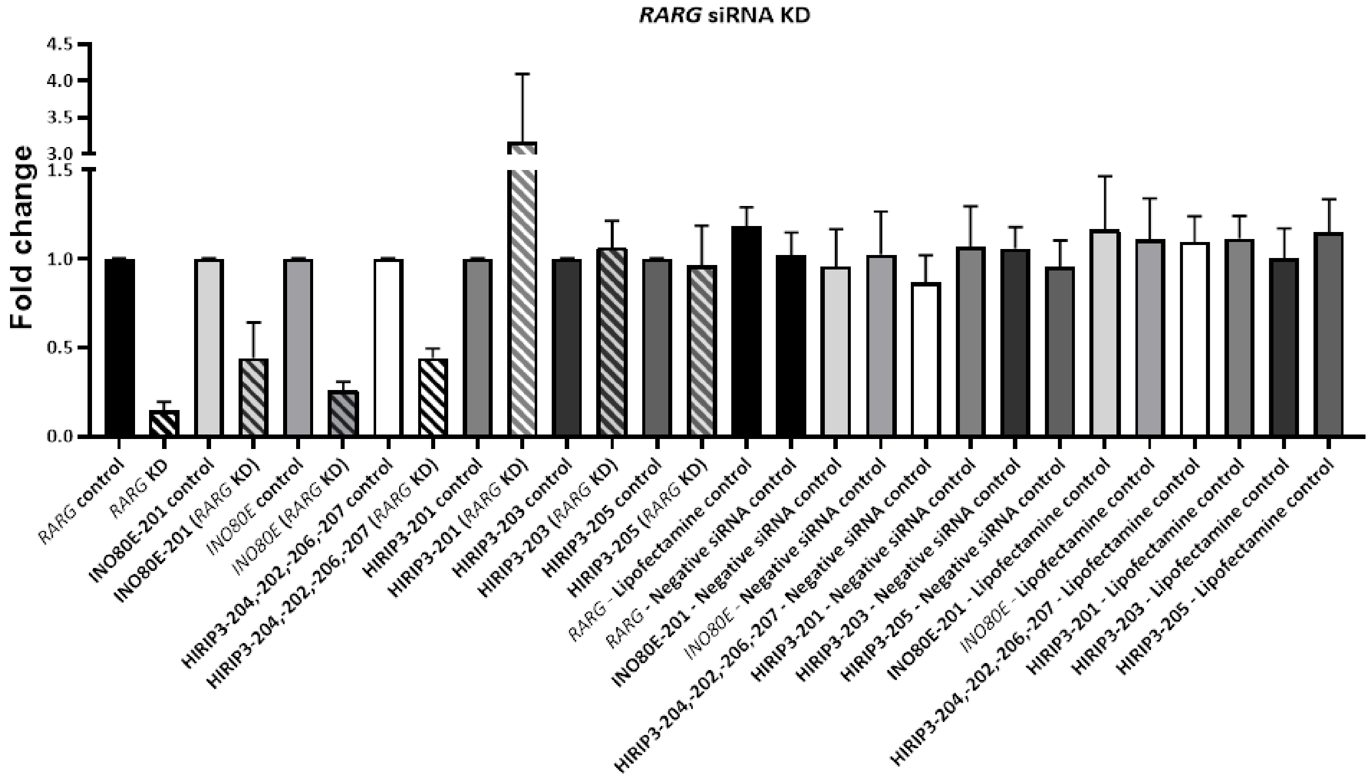

| **Brown-Forsythe and Welch ANOVA test; Dunnett's T3 multiple comparisons test** | **Summary** | **Adjusted P Value** |
|---|---|---|
| *RARG* control vs. *RARG* KD | **** | <0,0001 |
| *RARG* control vs. *RARG* - Lipofectamine control | ns | 0,0752 |
| *RARG* control vs. *RARG* - Negative siRNA control | ns | >0,9999 |
| INO80E-201 control vs. INO80E-201 (*RARG* KD) | ** | 0,0037 |
| INO80E-201 control vs. INO80E-201 - Lipofectamine control | ns | 0,9983 |
| INO80E-201 control vs. INO80E-201 - Negative siRNA control | ns | >0,9999 |
| *INO80E* control vs. *INO80E* (*RARG* KD) | **** | <0,0001 |
| *INO80E* control vs. *INO80E* - Lipofectamine control | ns | >0,9999 |
| *INO80E* control vs. *INO80E* - Negative siRNA control | ns | >0,9999 |
| HIRIP3-204,-202,-206,-207 control vs. HIRIP3-204,-202,-206,-207 (RARG KD) | **** | <0,0001 |
| HIRIP3-204,-202,-206,-207 control vs. HIRIP3-204,-202,-206,-207 - Lipofectamine control | ns | 0,9736 |
| HIRIP3-204,-202,-206,-207 control vs. HIRIP3-204,-202,-206,-207 - Negative siRNA control | ns | 0,8121 |
| HIRIP3-201 control vs. HIRIP3-201 (*RARG* KD) | * | 0,013 |
| HIRIP3-201 control vs. HIRIP3-201 - Lipofectamine control | ns | 0,7738 |
| HIRIP3-201 control vs. HIRIP3-201 - Negative siRNA control | ns | >0,9999 |
| HIRIP3-203 control vs. HIRIP3-203 (*RARG* KD) | ns | >0,9999 |
| HIRIP3-203 control vs. HIRIP3-203 - Lipofectamine control | ns | >0,9999 |
| HIRIP3-203 control vs. HIRIP3-203 - Negative siRNA control | ns | >0,9999 |
| HIRIP3-205 control vs. HIRIP3-205 (*RARG* KD) | ns | >0,9999 |
| HIRIP3-205 control vs. HIRIP3-205 - Lipofectamine control | ns | 0,9051 |
| HIRIP3-205 control vs. HIRIP3-205 - Negative siRNA control | ns | >0,9999 |

**Figure S8.** *RARG* gene knock-down effect on the *INO80E* and *HIRIP3* genes (ns – not significant, *- p<0.05; **- p<0.01; **** - p<0.0001).

**Table S1.** List of the oligonucleotides.

| Biotinylated probe | Sequence | |
|---|---|---|
| HIRIP3_probe | AGTCCGGCTCTGAAGCCTCCAGCCCAGACTACTTTGGACCCCCAGCAAAGAATGGGGTGGCAGCAGAAGTCAGCCCAGCCAAAGA | |
| CCT3_probe | GCGTGCCGCTGCAAACTGGCCTCTGGGCCGGGGGCGAGCAGCCCCCGGGAGGCCGAGTGCATCTGTTGGACCGTGCGAGGAGAAGAAAAA | |
| **Primers – qPCR** | | |
| **Primer name** | **Sequence** | **Product size** |
| qHIRIP3-205_F | GCGCATCTCTTATGTCCCCA | 138 bp |
| qHIRIP3-205_R | ACTTCGGATAGGTTGCCTCG | |
| qHIRIP3-201_F | CCGGAGAAAGTCAGCTAGGG | 149 bp |
| qHIRIP3-201_R | TCTGCACTGGGGGTTCTCTA | |
| qHIRIP3-203_F | CTGTTGGGCTCCTGTTGCTC | 148 bp |
| qHIRIP3-203_R | AGAAGCTGGTGATTCTCTCCTC | |
| qHIRIP3-204,202,206,207_F | TCTCGTCTGGCAAAGTTCTCG | 210 bp |
| qHIRIP3-204,202,206,207_R | CGGAAGAAGCTACGGGTGAA | |
| qINO80E_F | CCACTGCTTGGGGTAGCG | 128 bp |
| qINO80E_R | AGGAACTTGAGCTTCCGCTT | |
| qINO80E-201_F | AGCGTCCGCCATTTTTGT | 157 bp |
| qINO80E-201_R | TGTGCCGTCAGGACTACAAC | |
| qRARG_F | TGTGCGAAATGACCGGAACA | 237 bp |
| qRARG_R | ATGCACTTGGTAGCCAGCTC | |
| qGAPDH_F | GATGACAAGCTTCCCGTTCTC | 197 bp |
| qGAPDH_R | TGAAGGTCGGAGTCAACGGA | |
| qMALAT1_F | TTTTAGCAACGCAGAAGCCC | 166 bp |
| qMALAT1_R | ATACCACCACCTGGAATGGC | |
| qCCT3_F | CAAGACGGAGTTTCACCATGT | 163 bp |
| qCCT3_R | AAAGGAAGACAATGGAGGTGGT | |
| **Primers – ChIP-qPCR** | | |
| **Primer name** | **Sequence** | **Product size** |
| chINO80E_HIRIP3_F3 | CCTTTCCAACCCCATCTCCAAC | 173 bp |
| chINO80E_HIRIP3_R3 | CGAGCCGCTGTCAAACGC | |
| chINO80E_HIRIP3_F4 | CCGGGATTGACGGCTCCC | 175 bp |
| chINO80E_HIRIP3_R4 | CCAACTGCCCAGAGAAAACAAG | |
| chINO80E_HIRIP3_F5 | AGAACTTTGCCAGACGAGACT | 229 bp |
| chINO80E_HIRIP3_R5 | CTCGCGAGGATGCCTTTTCT | |
| chINO80E_HIRIP3_F7 | CAGCGTCCGCCATTTTTGTG | 157 bp |
| chINO80E_HIRIP3_R7 | TGTGCCGTCAGGACTACAAC | |
| chINO80E_HIRIP3_F8 | GGGAGTTGTAGTCCTGACGG | 193 bp |
| chINO80E_HIRIP3_R8 | AACTTGAGCTTCCGCTTCAGA | |
| chINO80E_HIRIP3_F9 | TACCGGAATCTGAAGCGGAA | 197 bp |
| chINO80E_HIRIP3_R9 | CTCCTGTATCACGGGCAAGG | |
| chINO80E_HIRIP3_F10 | CCAGTGCCCTTCTGTAGTGT | 152 bp |
| chINO80E_HIRIP3_R10 | TCAGACAACTTGGGTGTCCC | |
| chINO80E_HIRIP3_F11 | CATGCCTAGCGCTTCCAGTTC | 179 bp |
| chINO80E_HIRIP3_R11 | TCTTTCCTTGCTAGGCTGGCTC | |
| **Primers – R-loops detection, RNAseH1 overexpression** | | |
| **Primer name** | **Sequence** | **Product size** |
| rINO80E_MET_F | CCGGGATTGACGGCTCCC | 277 bp |
| rINO80E_MET_R | AACTAATTAAATTTCACCCAAAAAATAC | |
| dINO80E_MET2_F2 | GGGGATAAATTTTTTTGTGGTTTT | 480 bp |
| dINO80E_MET2_R2 | CCACCCACTTTACACCTAAATAAACTA | |
| qRNAseH1_F | AGCCCGGAAGTTTCAGAAGG | 126 bp |
| qRNAseH1_R | GTGCTTTGCATACGGCTCTG | |

| **gRNAs** | |
|---|---|
| **Primer name** | **Sequence** |
| gINO80E_MET2_F1 | CACCCTGTCCCAGTGTCCCCCGGG |
| gINO80E_MET2_R1 | AAACCCCGGGGGACACTGGGACAG |
| gINO80E_MET2_F2 | CACCCTACGCCATCCCCAGCGCAG |
| gINO80E_MET2_R2 | AAACCTGCGCTGGGGATGGCGTAG |
| gINO80E_MET2_F3 | CACCGCCGGGAGTTGTAGTCCTGA |
| gINO80E_MET2_R3 | AAACTCAGGACTACAACTCCCGGC |
| gINO80E_MET2_F4 | CACCTGCAGCGTCTCCGGAAGTGG |
| gINO80E_MET2_R4 | AAACCCACTTCCGGAGACGCTGCA |
| gINO80E_MET2_F5 | CACCGCCGGTCATGAACGGGCCGG |
| gINO80E_MET2_R5 | AAACCCGGCCCGTTCATGACCGGC |
